# Single-cell atlas of the fetal Down syndrome cortex uncovers chromosome 21 regulators of intellectual disability gene networks

**DOI:** 10.64898/2025.12.29.696602

**Authors:** Michael Lattke, Wee Leng Tan, Salil Kalarikkal Sukumaran, Kagistia Hana Utami, Marcos Sintes, Srinivasan Sakthivel, Jonathan Tan, Auriel Lim, Bansal Aysha Vibhavari, Katerina Rekopoulou, Nik Matthews, Ivan Alic, Željka Krsnik, Dean Nizetic, Boaz P. Levi, Vincenzo De Paola

## Abstract

Down syndrome (DS), caused by trisomy of chromosome 21, is the leading genetic cause of intellectual disability, yet the mechanisms disrupting fetal brain development remain unclear. We performed single-cell transcriptomic and chromatin accessibility profiling of approximately 250,000 cells from 15 DS and 15 control human fetal cortices (10–20 weeks post-conception). Our analysis revealed a subtype-specific reduction in RORB/FOXP1-expressing excitatory neurons and widespread disruption of neurodevelopmental transcriptional programs. Chromosome 21 transcription factors BACH1, PKNOX1, and GABPA emerged as dosage-sensitive hubs regulating genes linked to intellectual disability. Antisense oligonucleotide-mediated normalization of these factors in human neural progenitors *in vitro* partially rescued target gene expression. Benchmarking a humanized *in vivo* model captured additional molecular and cellular signatures of DS, complementing the *in vitro* model. Together, this resource defines the gene-regulatory landscape underlying cortical development in DS and highlights candidate molecular targets and preclinical models for future intervention studies.

**Highlights:**

- Single-cell atlas of Down syndrome fetal cortex links transcriptional dysregulation to reduction of layer 4 neurons
- Chromosome 21 transcription factors PKNOX1, BACH1, and GABPA drive intellectual disability gene dysregulation
- Transplanted human cells model late-stage DS phenotypes, bypassing scarcity of fetal tissue
- ASO targeting chromosome 21 transcription factors restores DS-associated molecular signatures

## Main

Trisomy (Ts21) of chromosome 21 (Chr21) is the most common chromosomal abnormality in humans and a major cause of intellectual disability^1–3^. Despite the severe impact on quality of life, there are no effective treatments available for the intellectual disability and other neurological manifestations of DS, such as early-onset Alzheimer’s-like dementia, behavioral impairments, and infant seizure susceptibility^1–3^. Imaging and postmortem studies have revealed reduced fetal brain volume starting from gestational week 23 (GW23; i.e. post-conceptional week (PCW) 21), linked to decreased neural progenitor proliferation and excitatory neuron production from GW18, increased astrogliogenesis, and impaired dendrite and synapse formation in postnatal stages (reviewed in^2^). Traditional candidate-based genetic approaches in mice and stem cell models have identified various genes that may contribute to these phenotypes, such as DYRK1A, DSCAM, OLIG2, APP, and IFNAR1/2 (for review, see^2,4,5^). Previous efforts to comprehensively understand global gene expression dysregulation in the DS brain using stem cells, mouse models and post-mortem human samples have provided valuable insights into altered molecular pathways in Ts21^6–10^. However, *post-mortem* human studies have either focused on adult stages^11,12^, or lacked cellular resolution^13^. As a result, despite significant insights from previous work, it is still unclear how the subtle (approximately 1.5-fold) increase in gene dosage of ∼ 200 Chr21 genes due to Ts21 impacts the development and function of various cell populations in the human cortex, the region central to cognitive functions. Specifically, we lack insights into how these changes affect the genomic programs that govern human cortical development and function during the critical period (∼ PCW 10-20) when most neurons are generated^14^. This period lays the foundation for cortical organization and connectivity, likely contributing to the neurological features observed in DS. Here we combined multimodal network analysis of single-nucleus RNA sequencing (snRNA-seq) and single-cell assay for transposase-accessible chromatin sequencing (scATAC-seq) from the same cells, with human neuron *in vitro* and *in vivo* modelling to determine how increased Chr21 gene dosage impacts the development and function of various human cortical cell populations. We further defined the earliest cellular and molecular perturbations in DS cortical development, as well as the underlying mechanisms, creating a publicly available, cellular-resolution atlas of gene expression, chromatin accessibility and the regulatory architecture of the human DS cerebral cortex at PCW 10-20.

## Results

### A single-cell gene expression and chromatin accessibility atlas of the human fetal cortex in Down syndrome

To identify early molecular and cellular changes in the developing DS cortex, we performed combined single-cell transcriptional and chromatin accessibility profiling (10X Genomics multiome technology) of human brain samples from fetuses with Ts21/DS, and diploid controls (CON) from elective terminations of pregnancy, i.e. likely free of developmental defects. We acquired 20 CON and 19 DS fetal brain samples from surgical terminations (see Methods, Supplementary Table 1) spanning PCW 10 to 20, the period from early cortical neurogenesis to early gliogenesis (Fig. 1a-b). With immunostainings we identified samples containing both the ventricular zone (VZ), with PAX6-expressing cortical progenitors, and the cortical plate (CP), which included newly generated neurons expressing CTIP2 (BCL11B) and SATB2, markers for cortical layer (L)5/6 and L2/3 excitatory neurons, respectively (Extended Data Fig. 1a). After excluding two poorly preserved samples, we performed single nucleus RNA/ATAC sequencing. We performed stringent quality controls with the obtained datasets, removing low quality samples and cells, as well as 3 samples mapping to non-cortical regions of a reference atlas^15^ (see Methods, Extended Data Fig. 1, Extended Data Table 1), retaining 248,998 high-quality cells from 15 CON and 15 DS samples. We integrated these samples, performed dimensionality reduction by Uniform Manifold Approximation and Projection (UMAP) and clustering of transcriptionally similar cells into 21 populations (see Methods). We then characterized these populations, which mapped reproducibly between samples, using established marker genes and the reference atlas^15^ (Fig. 1c-d; Extended Data Fig. 1c-d, 2a-b). Most cells expressed high levels of neural lineage markers, including four populations expressing the radial glia/astrocyte markers SLC1A3 (also known as GLAST), SOX9, PAX6, NES and HOPX (RG_c0/c11, RG_prol_c8, AST_c13). One of these populations (RG_prol_c8) co-expressed proliferation markers MKI67, TOP2A, CDK1, and one (AST_c13) co-expressed astrocyte-specific markers ALDH1L1, GFAP, GJA1 (CX43), AQP4 and S100B. Three populations expressed excitatory intermediate progenitor cell markers EOMES, GADD45G, ASCL1, including one co-expressing proliferation markers (IPC_c5/c12, IPC_prol_c9). Seven populations expressed excitatory cortical pyramidal neuron markers NEUROD2 and NEUROD6, including one (NEU_TLE4_c3) expressing the L5/6 neuron markers TLE4, TBR1 and BCL11B (CTIP2), one expressing the L4 markers RORB and FOXP1 (NEU_RORB_c4), three expressing the L2/3 marker CUX2 (NEU_CUX2_c0/c2/c10), and two minor populations only lowly expressing subtype markers (NEU_low_c17/c20). Consistent with their putative identity as cortical excitatory lineage cells, these clusters expressed the dorsal forebrain marker FOXG1. Four populations expressed GAD1, GAD2, and markers of ventrally, ganglionic eminences-derived cells (LHX6, DLX2, ADARB2), consistent with GABAergic interneuron identity. These also expressed different interneuron subtype markers, including SST, CALB2 (Calretinin) and RELN (NEU_SST_c6, NEU_CALB_c7, NEU_RELN_c14/c15). We also found three minor populations expressing the oligodendrocyte lineage markers PDGFRA, CSPG4 (NG2), MBP and MOG, putative oligodendrocyte precursor cells (OPC_c16), microglia markers (CX3CR1, ITGAM; MIC_c19), or endothelial cell and pericyte markers PECAM1, CLDN5, MCAM and PDGFRB, putative vascular cells (VASC_c18).

**Figure 1.**
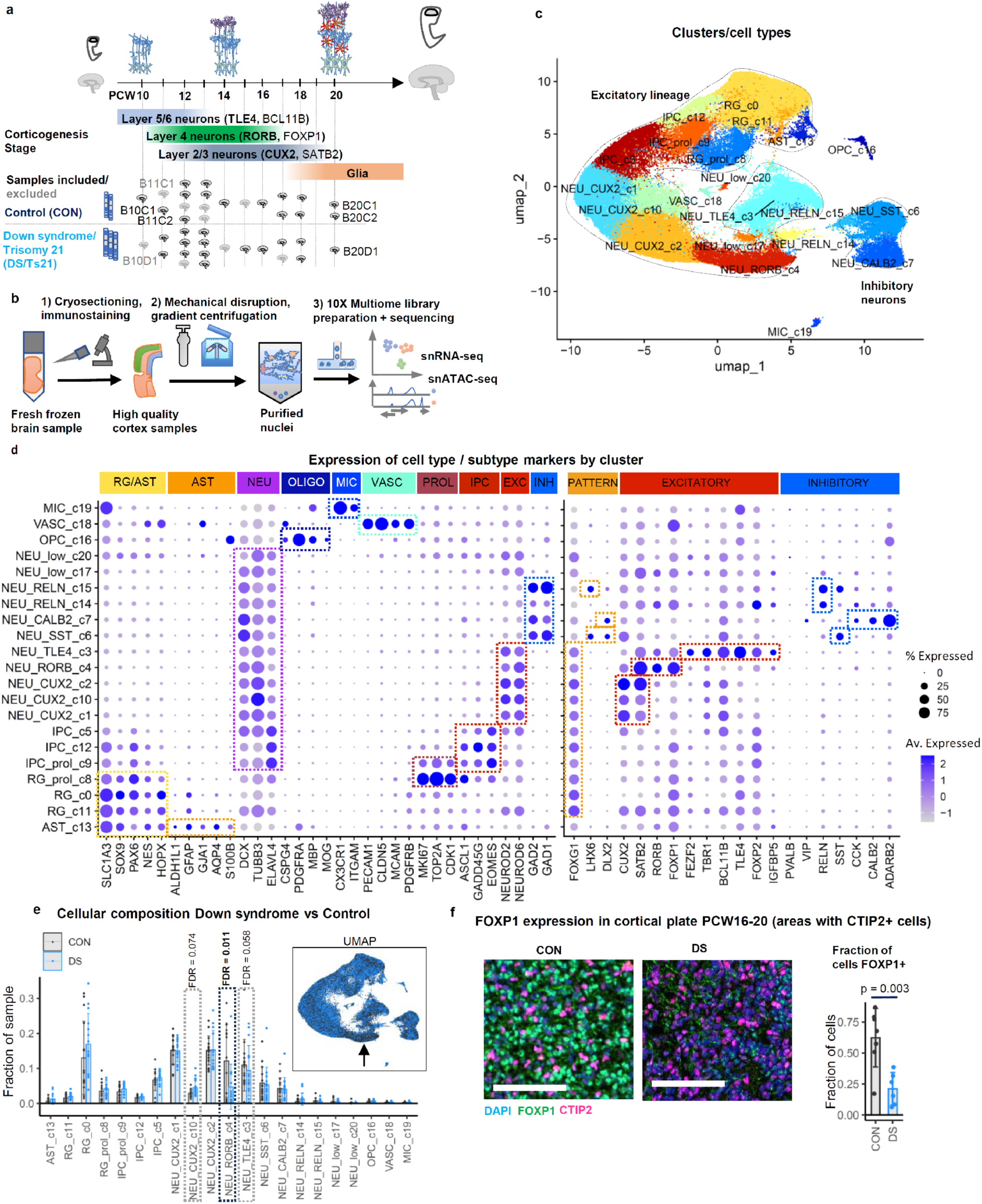
A single-cell gene expression and chromatin accessibility atlas of the human fetal cortex in Down syndrome. **a**, Stages of cortical development covered by fetal tissue samples used for this dataset. **b**, Experimental pipeline for processing of brain samples for control stainings, nuclei extraction and combined single cell transcriptome and chromatin accessibility analyses. **c**, Cell type assignment of identified cell clusters (UMAP plot). **d**, Expression of marker genes used to assign clusters to cell types (left) and subtypes (right). **e**, Abundance of cell populations in control (CON) and Down syndrome (DS) samples. Barplot showing individual samples with False Discovery Rate (FDR) for DS vs CON from sccomp compositional analysis^18^ (other clusters FDR>0.05); Inlay: combined UMAP plot (arrow: NEU_RORB_c4 cluster reduced in DS). **f**, FOXP1 immunostaining in CON and DS brains from PCW16-20; Sections from cryopreserved brains used for sequencing analyses (representative CTIP2 positive cortical plate areas from images analyzed for quantification); scale bar: 100 μm; statistical analysis: 2-tailed t-test. See also Extended Data Fig. 1 and 2, Supplementary Table 1.

**Extended Data Fig. 1.**
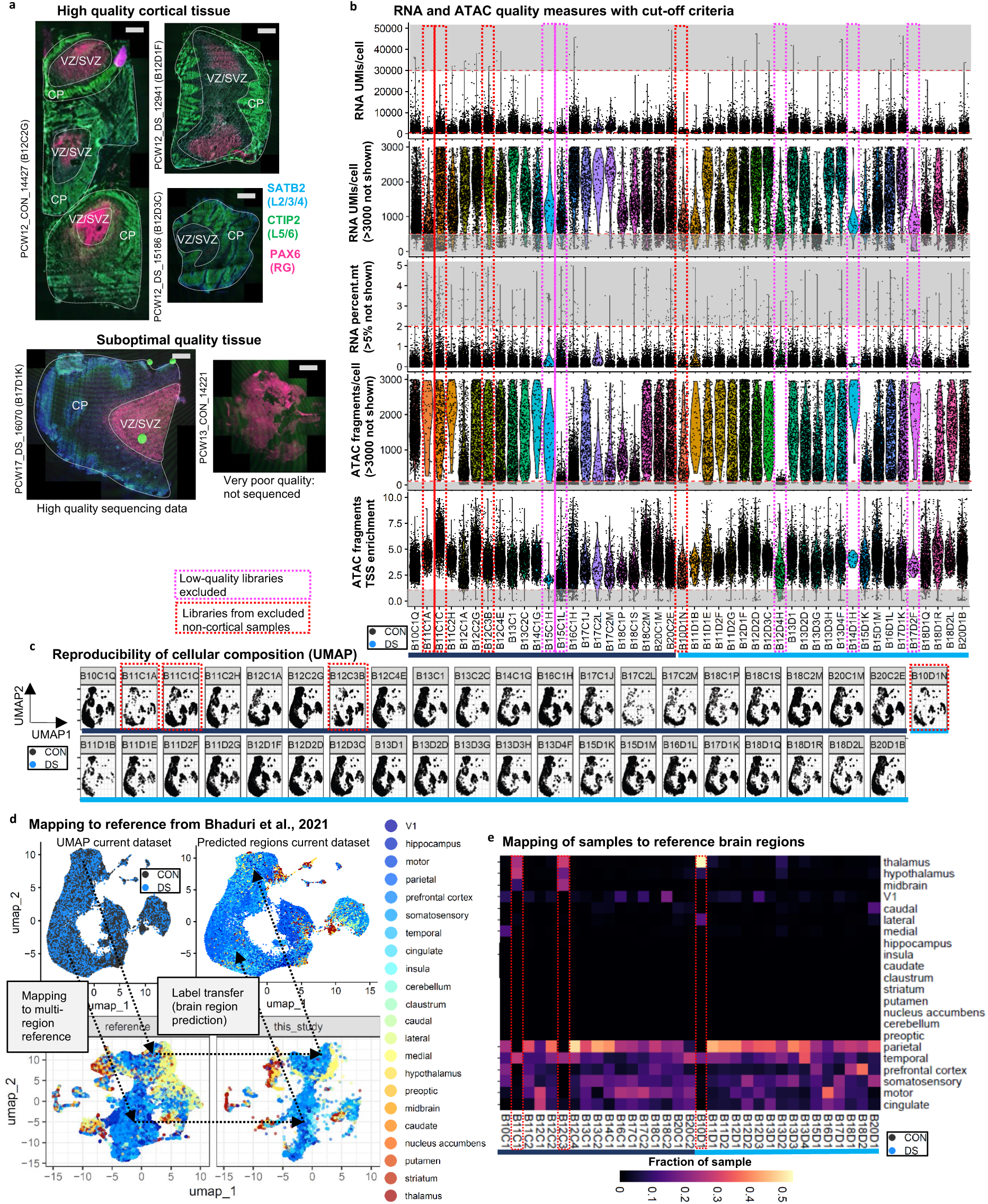
Quality control and identification of non-cortical samples in the fetal brain single-cell multiomic atlas. **a**, Immunostaining for markers of the cortical progenitor zone (PAX6) and cortical plate neurons (SATB2, CTIP2) to identify samples suitable for single-cell multiome analysis. Representative confocal images, scale bar: 1 mm. **b**, RNA- and ATAC-seq quality measures per cell by library; grey shadows delimited by the red dotted cut off lines: excluded low-quality cells; purple dotted boxes: libraries excluded due to high fraction of low-quality cells; red dotted boxes: libraries from excluded non cortical samples. **c**, UMAP plot split by libraries included in the dataset. Note reproducible contribution of libraries/samples to cell populations. **d-e**, Regional identity predicted by mapping of cells / samples to multi-region reference dataset^15^; highlighted by dotted green boxes (e): Excluded samples with predicted non-cortical cells.

**Extended Data Fig. 2.**
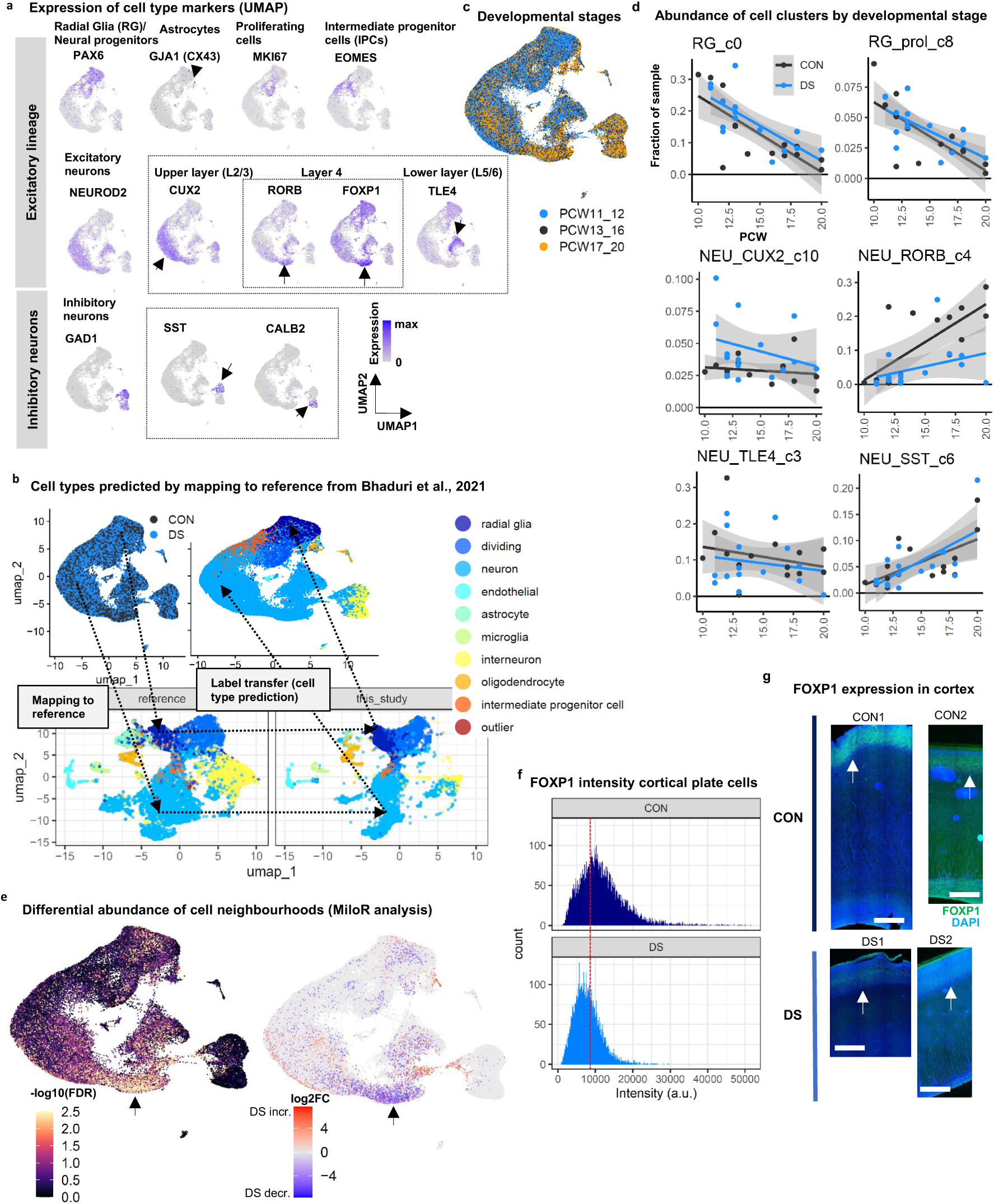
Characterization of cell populations in the fetal brain single-cell multiomic atlas. **a**, Cell type marker gene expression projected on UMAP dimensionality reduction plot. Arrows: small populations with high marker expression; rectangles: subtype markers. **b**, Cell-type predictions from mapping to reference dataset15. **c**, Contribution of samples from different stages to cell populations (UMAP plot). **d**, Changes of cell proportions in samples over developmental time. **e**, Differential abundance of cells in local neighborhoods quantified with MiloR^17^. Negative log10-transformed adjusted p-value (left) and log2-transformed fold change by neighborhood (right); Arrows: population of RORB/FOXP1-expressing neurons. **f-g**, FOXP1 immunostaining in CON and DS brains from PCW16-20; Intensity distribution of FOXP1 staining of cryosections (f) with cut-off for quantification (red line); Sections from two paraffin-embedded CON and two DS brains (g, arrow: cortical plate); scale bars: 1 mm.

Most populations were present in all samples (Figure 1e), although their abundance varied strongly between samples. Nevertheless, compositional changes broadly followed expected developmental patterns, with later stages including more late-developing RORB/FOXP1-expressing neurons and SST-interneurons (NEU_RORB, NEU_SST), and fewer progenitors (Extended Data Fig. 2c-d).

Notably, L4-like neurons expressing RORB/FOXP1 were dramatically reduced in DS samples, particularly at later stages (Fig. 1e, Extended Data Fig. 2d), a defect previously reported only in adult DS^11^ and in Alzheimer’s disease (AD) patients^16^. We confirmed this phenotype with an alternative cluster-free analysis (MiloR^17^), and FOXP1 immunostainings of tissue from PCW16-20, including in two additional pairs of well-preserved paraffin-embedded brains and sections from the cryopreserved brains used for the transcriptomic analyses (Methods, Fig. 1f, Extended Data Fig. 2e-g). Contrary to previous reports of reduced proliferating progenitors and increased interneuron or astrocyte numbers in DS at later stages^2^, we found no changes in progenitor, interneuron or astrocyte numbers at PCW10-20 (Fig. 1e).

Overall, our mid-gestation dataset primarily encompasses neural cells, including the entire excitatory lineage, multiple interneuron populations, and early glial cells. It reveals a marked reduction in putative L4 pyramidal excitatory neurons expressing RORB/FOXP1 as earliest cellular phenotype, while other previously reported compositional changes could not be detected, suggesting they may arise at a later stage.

### Gene expression changes mainly affect excitatory neurons and are linked to neural development and function

Next, we investigated how Ts21 affects global gene expression, to identify cell types and genetic programs that may contribute to the biological features associated with DS. We compared gene expression between DS and CON for each cell cluster with a pseudobulk-based approach with low differential expression threshold (1.2-fold, see Methods), to allow detection of subtly deregulated genes, including Chr21 genes, expected to be upregulated 1.5-fold on average due to the presence of the additional copy of Chr21.

As expected^19^, the 87 differentially expressed Chr21 genes were exclusively upregulated, in a wide range of cells (Extended Data Fig. 3a, Extended Data Table 2). Of the remaining 732 differentially expressed genes, the majority were identified in RORB/FOXP1-expressing neurons (NEU_RORB_c4)—whose abundance is reduced—as well as in TLE4-expressing neurons (NEU_TLE4_c3), and in two smaller populations (NEU_RELN_c14, NEU_low_c17). To identify biological processes likely affected by the observed transcriptomic changes, we performed a Gene Ontology (GO) analysis (Extended Data Fig. 3b, Supplementary Table 2). The most prominent among the 114 enriched GO terms were terms related to neurodevelopmental processes, whose deregulation could contribute to cognitive impairment in DS, such as ‘*forebrain development’, ‘neural precursor cell proliferation’*, ‘*regulation of neuron differentiation*’, ‘*axonogenesis’*, or ‘*dendrite development*’.

**Extended data Fig. 3.**
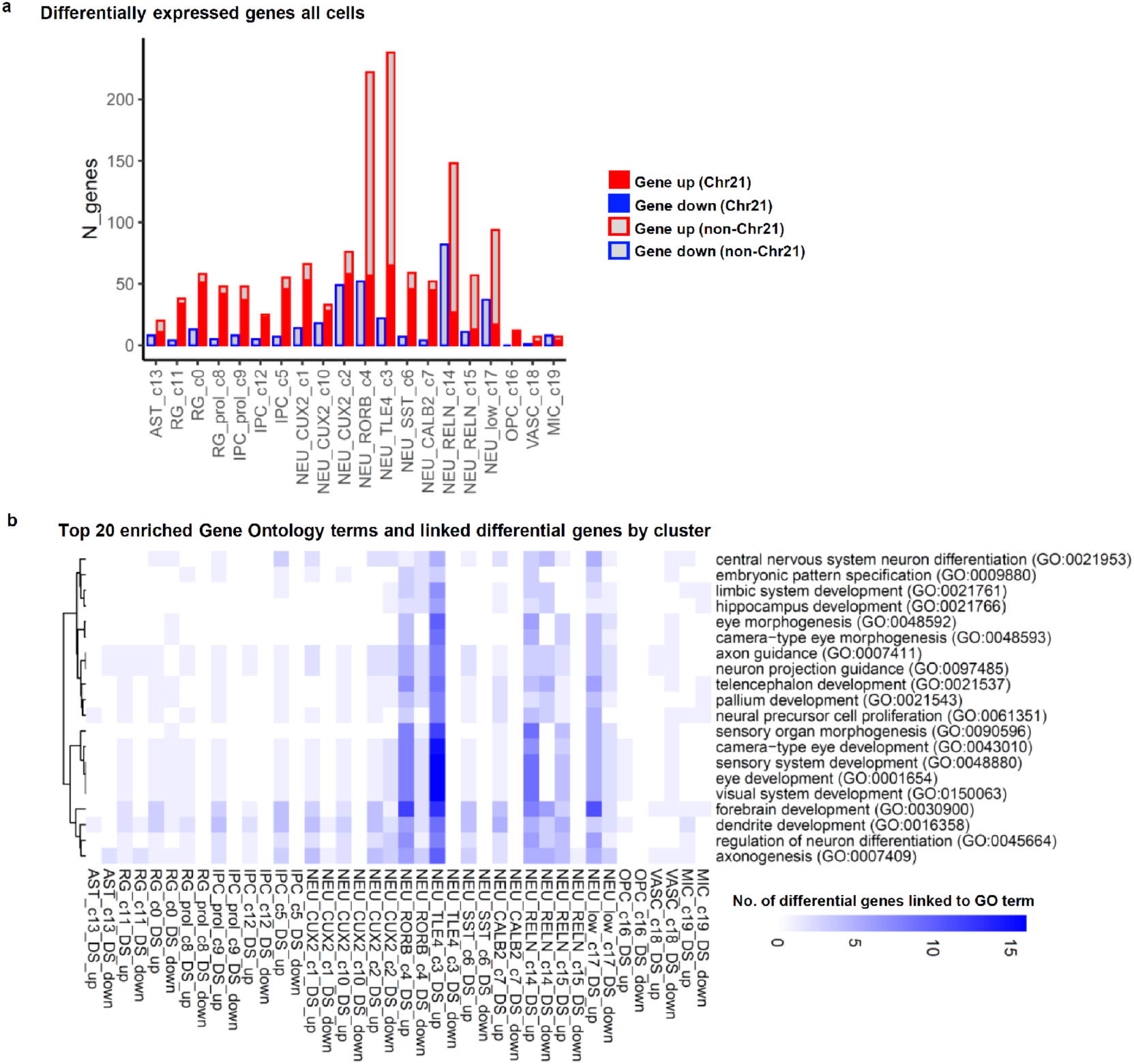
Expression changes in the complete dataset comprising all cell populations and gene ontology (GO) terms enriched for differential genes. **a**, Number of genes differentially expressed between DS and CON samples by cluster; DESeq2 pseudobulk analysis with Wald test, threshold padj<0.10, log2FoldChange > log2(1.2). **b**, Biological processes linked to differentially expressed genes: heatmap showing number of differentially expressed genes in Top20 enriched gene ontology (GO) terms by cell cluster.

As most inhibitory neurons and microglia — cell types previously reported to be affected in DS^11^— showed only limited transcriptional changes in our dataset, we focused our analyses on excitatory neurons, where we detected substantially more widespread transcriptional alterations. To investigate these changes in more detail, we subsetted and re-clustered our dataset, retaining only excitatory neurons and their progenitors, including putative immature astrocytes, which derive from the same progenitors and are not unambiguously distinguishable from radial glia (Methods, Fig. 2a, Extended Data Fig. 4a-b). In all cell clusters in this subset, together we detected 672 differentially expressed genes (Fig. 2b-c, Supplementary Table 3), including many non-Chr21 genes in RORB/FOXP1 and TLE4 expressing neurons (NEU_RORB_s4, NEU_TLE4_s3; Fig. 2b), which are also reduced in abundance (Extended Data Fig. 4c). Differential genes were enriched for GO terms linked to processes impaired in DS, including ‘*cognition*’ and neurodevelopmental terms, such as ‘*forebrain development’, ‘sensory organ morphogenesis’*, ‘*axonogenesis’*, ‘*dendrite development*’, ‘*gliogenesis*’ and ‘*neural precursor cell proliferation’* (Fig. 2c, Supplementary Table 3)^2^. Genes showed generally only subtle changes and included Chr21 genes such as APP, GART and C21ORF91, as well as key transcription factors, receptors and ligands involved in cortical neuron specification and differentiation located on other chromosomes, such as NEUROG2, FEZF2, FOXP1, NTRK2, FGFR2, NOTCH1 and WNT4 (Fig. 2d). Of note, FOXP1, whose mutations can cause intellectual disability, and which has been implicated in impaired generation of L4 to L6 excitatory neurons expressing RORB or TLE4, respectively^20^, also showed reduced nuclear immunoreactivity in tissue sections (Fig. 1f; Extended Data Fig. 2f-g), validating downregulation on protein level.

**Fig. 2.**
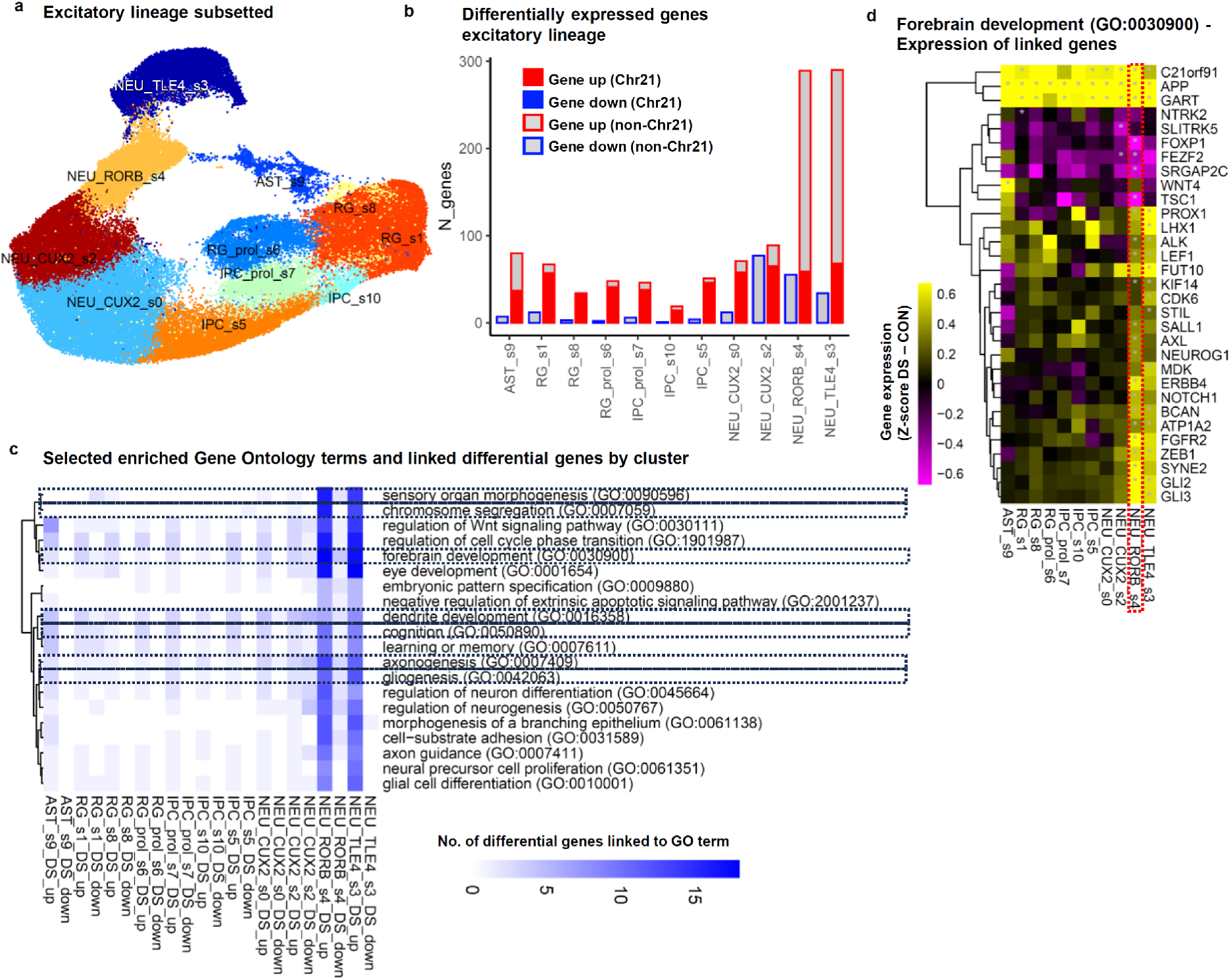
Gene expression changes mainly affect excitatory neurons and are linked to neural development and function. **a**, Cell populations of the subsetted and re-clustered excitatory lineage PCW10-20 dataset (UMAP plot). Marker expression for cluster assignment see Extended Data Fig. 4a-b. **b**, Number of genes differentially expressed between DS and CON samples by cluster; DESeq2 pseudobulk analysis with Wald test, threshold padj<0.10, log2FoldChange > log2(1.2). **c**, Biological processes linked to differentially expressed genes: heatmap showing number of differentially expressed genes in selected enriched gene ontology (GO) terms by cell cluster; highlighted: GO terms referred to in main text. **d**, Differentially expressed genes linked to GO term ‘*forebrain development*’ by cluster; heatmap colored by relative expression in DS vs CON (difference vst-normalized gene expression z-score DS vs CON samples). Highlighted by red dotted box: RORB/FOXP1-expressing neurons reduced in DS; Grey asterisks indicate padj<0.10. See also Extended Data Fig. 3-6, Supplementary Tables 2, 3.

**Extended data Fig. 4.**
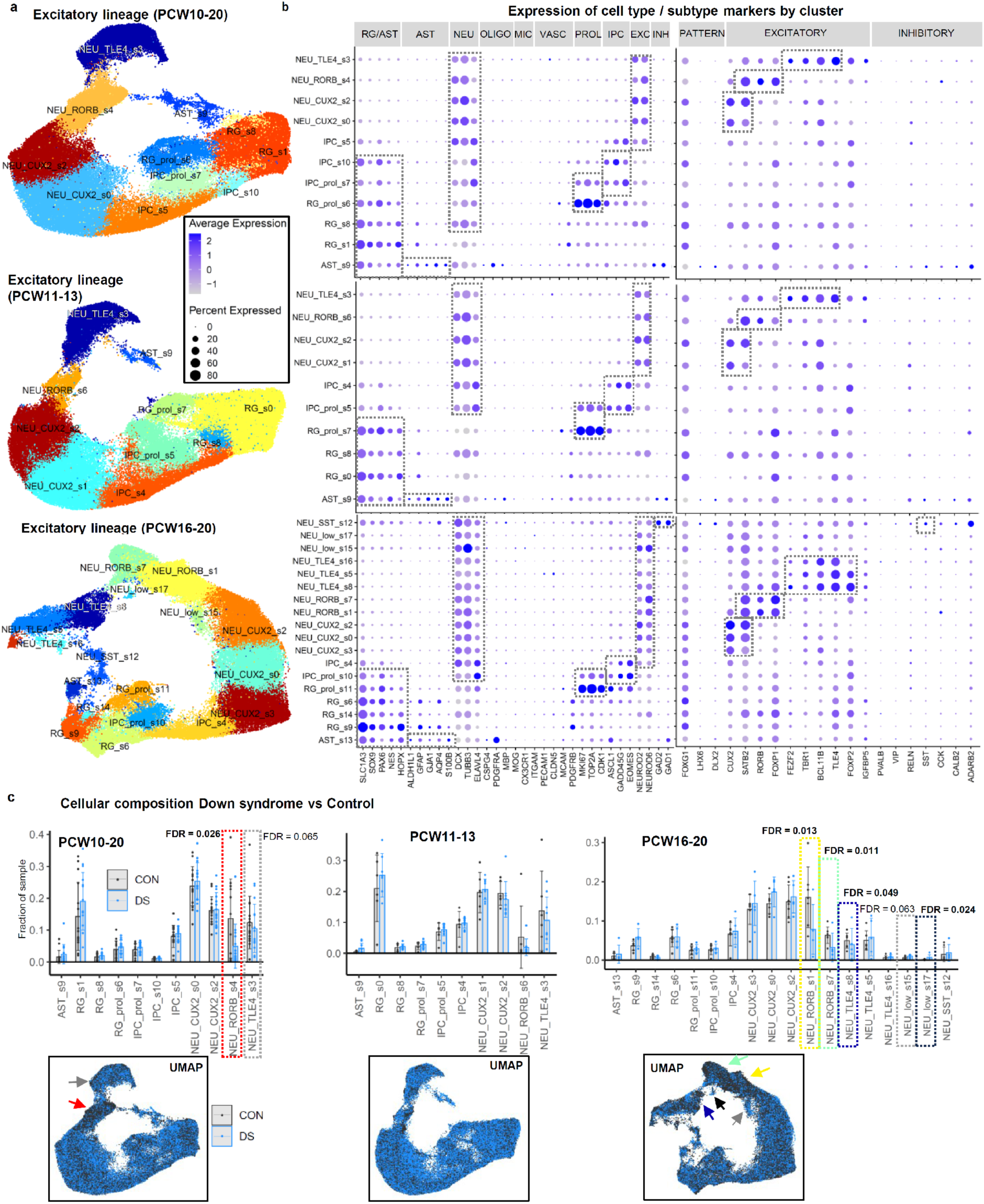
Subsetting of excitatory lineage cells from complete dataset (PCW10-20) or samples from early (PCW11 to PCW13) or late stages (PCW16 to PCW20) separately. **a**, Dimensionality reduction, re-clustering and cell type assignment of identified excitatory lineage cell clusters (UMAP plot). **b**, Expression of marker genes used to assign clusters to cell types/subtypes. **c**, Abundance of cell populations in control (CON) and Down syndrome (DS) samples. Barplot showing individual samples with False Discovery Rate (FDR) for DS vs CON from sccomp compositional analysis^18^ (other clusters FDR>0.10) and combined UMAP plot (dotted boxes / arrows: clusters altered in DS, sccomp FDR<0.10).

Altered expression of the majority of these genes was confirmed with Nebula, an alternative cell-level differential expression analysis approach^21^, and recapitulated in published bulk-RNA-seq data from adult cortex and iPSC-derived neurons^12^ (Extended Data Fig. 5, Supplementary Table 3).

**Extended data Fig. 5.**
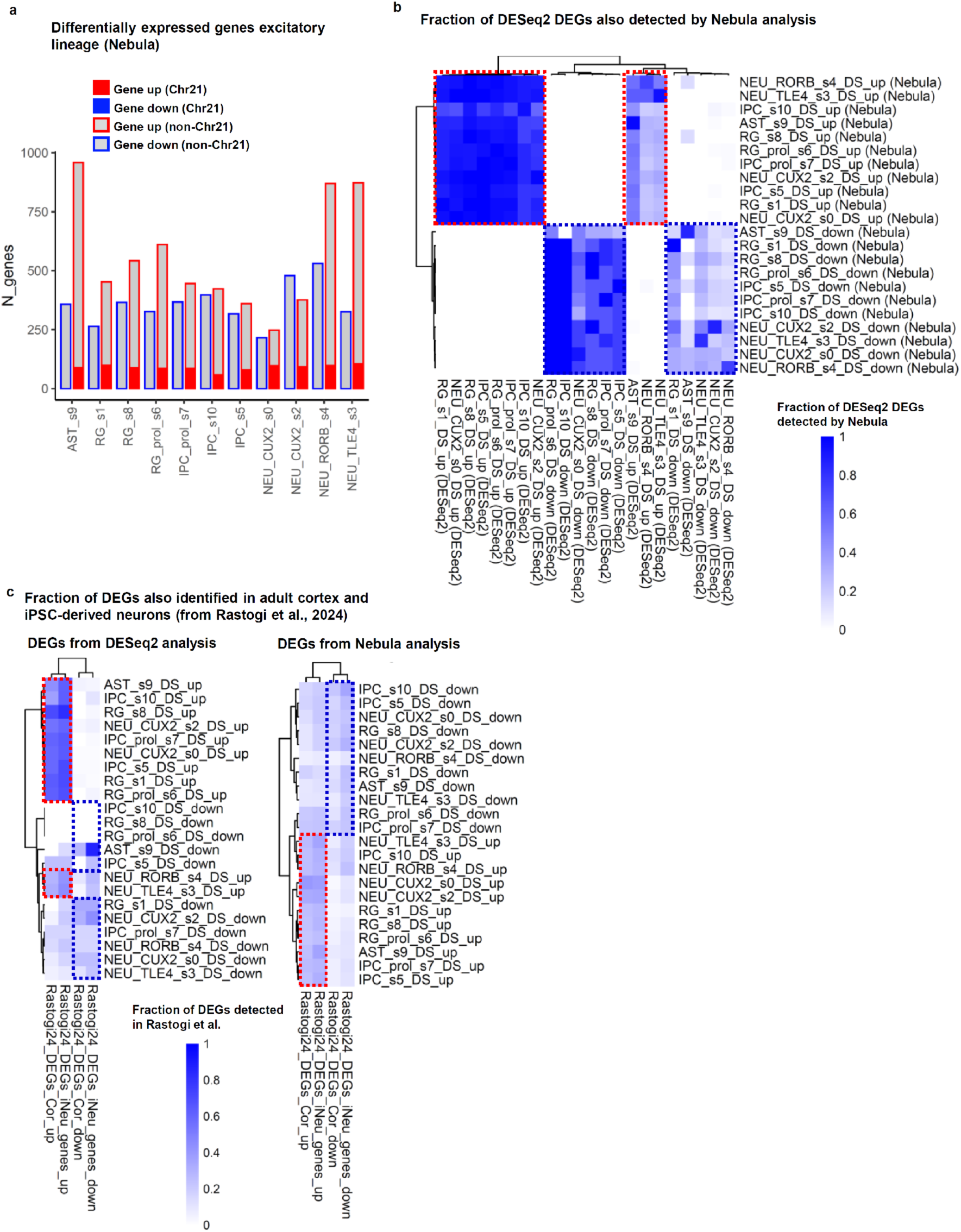
Validation of differentially expressed genes with alternative analysis approach and published data. **a**, Number of differentially expressed genes between DS and CON samples by cluster detected by Nebula analysis^21^; Threshold padj<0.05, log2FoldChange > log2(1.2). **b**, Fraction of differential genes per cluster identified with DESeq2 also detected by Nebula. **c**, Fraction of differential genes (identified with DESEq2 or Nebula) also detected in adult cortical tissue or iPSC-derived neurons (bulk-RNA-seq from^12^); note that Nebula identified more differential genes (a, see also Fig. 2b), but showed lower concordance with the published data (c). Boxes (b, c): Genes consistently upregulated (red) / downregulated (blue) in both analyses.

To pinpoint the timing of these changes, we separately subsetted excitatory cells from early (PCW11-13) and late-stage samples (PCW16-20; Extended Data Fig. 4, 6, Supplementary Table 3). Although a trend towards reduced RORB/FOXP1 expressing neurons was present already at PCW11-13, significant reductions of RORB/FOXP1 and TLE4 expressing neurons, were only observed at PCW16-20 (Extended Data Fig. 4c). 472 genes were already differentially expressed at PCW11-13 (Extended Data Fig. 6a-b, Supplementary Table 3), including 75 Chr21 genes and genes linked to neurodevelopmental programs (GO terms ‘*axonogenesis’*, ‘*dendrite development*’), and excitatory synaptic signaling (GO terms ‘*regulation of trans-synaptic signalling*’, ‘*ionotropic glutamate receptor signaling pathway*’).

**Extended data Fig. 6.**
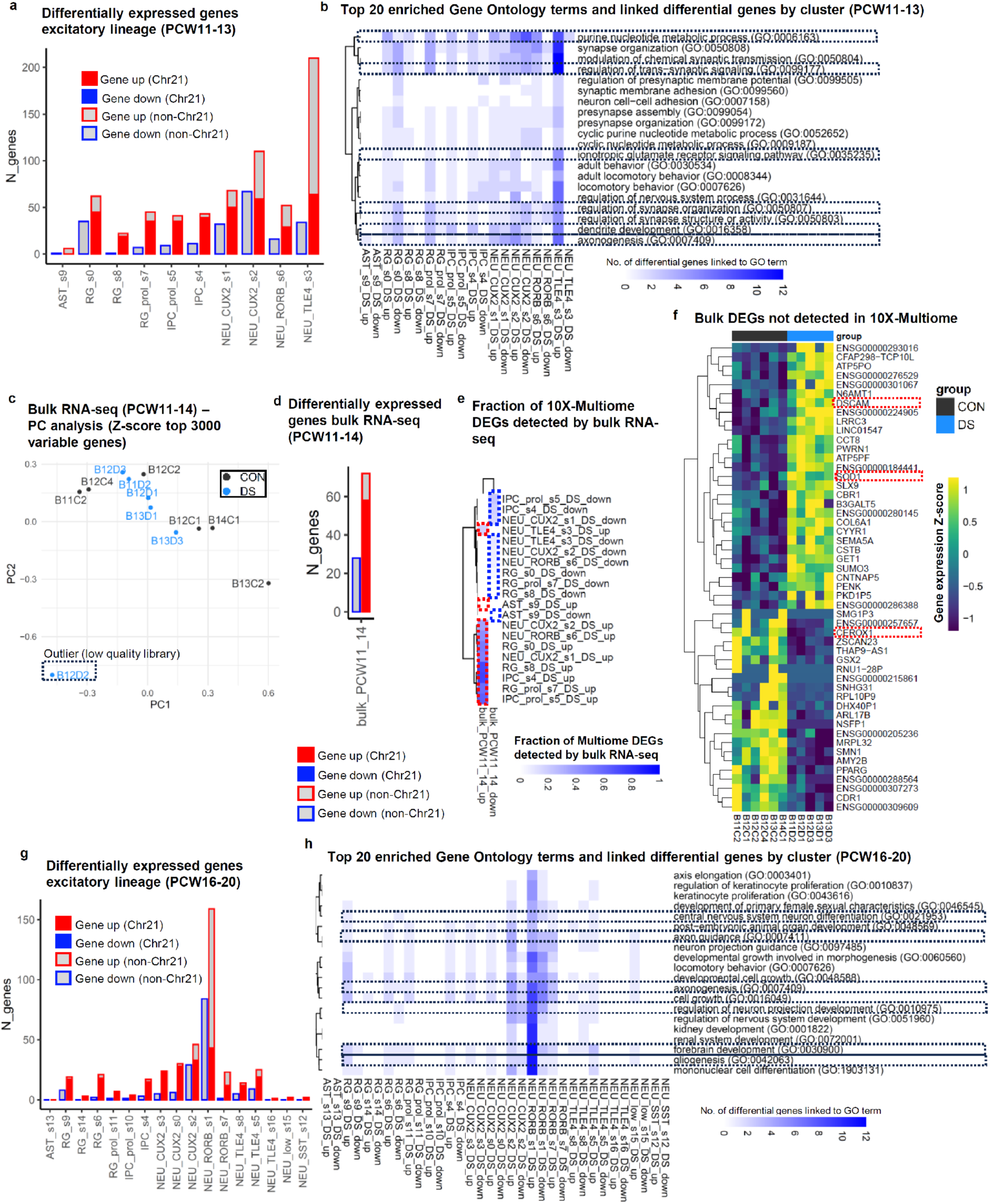
Expression changes in early and late excitatory lineage (PCW11-13 / PCW16-20). **a**, Number of genes differentially expressed between DS and CON samples PCW11-13 by cluster; DESeq2 pseudobulk analysis with Wald test, threshold padj<0.10, log2FoldChange > log2(1.2). **b**, Biological processes linked to differentially expressed genes PCW11-13: heatmap showing number of differentially expressed genes in Top20 enriched gene ontology (GO) terms by cell cluster. **c-f**, Complementary bulk-RNA-seq analysis from samples PCW11-14; Principal component analysis showing outlier low-quality library omitted from following analyses (c); Number of differential genes (d, DESeq2 pseudobulk analysis with Wald test, threshold padj<0.10, log2FoldChange > log2(1.2)); Fraction of differential genes in PCW11-13 populations also detected by bulk RNA-seq (e). Heatmap of expression of differential genes detected by bulk-RNA-seq, but not in single-cell analyses (f); Red dotted boxes: genes discussed in main text. **g, h**, Number of differential genes and enriched GO terms by cluster for PCW16-20 (as in a-b).

To further validate and characterize these early dysregulated programs, we performed bulk RNA-seq on a representative subset of samples from PCW11–14 (Extended Data Fig. 6c-f, Supplementary Table 3). This analysis confirmed that an average of 27.5% of genes significantly altered in the snRNA-seq analysis show concordant changes in bulk RNA, consistent with previous work^22^. This increases to an average of 46.8% for upregulated genes, reaching over 70% in two clusters. Importantly, the analysis also revealed additional differentially expressed genes not previously detected (Extended Data Fig. 6c-f, Supplementary Table 3). These included Chr21 genes implicated in DS-related phenotypes – such as DSCAM, SOD1^8,23^, as well as several largely uncharacterized non-coding RNAs, including CEROX1, which has been reported to regulate mitochondrial activity^24^—a pathway known to be altered in DS^25^.

In the PCW16–20 samples, 307 differential genes were detected (including 54 on Chr21), predominantly within the major RORB/FOXP1-expressing neuronal population (NEU_RORB_s1), consistent with greater heterogeneity at later developmental stages. Genes were enriched for GO terms such as ‘*forebrain development*’, ‘*regulation of neuron projection development*’ and ‘*gliogenesis*’ (Extended Data Fig. 6g-h, Supplementary Table 3).

Overall, this indicates that before PCW11–13, Ts21 perturbs transcriptional programs regulating excitatory neuron development and function. This leads to a selective deficit of RORB/FOXP1-expressing subtypes that may stem from impaired generation, maturation, or increased vulnerability^16^, and contribute to later neurological phenotypes in DS.

### Integrated gene regulatory network analysis predicts key mediators contributing to the deregulation of transcriptional programs downstream of Chr21 genes

We next asked how the increased gene dosage of Chr21 genes might cause the observed transcriptional alterations. Cell-type-specific accessibility of cis-regulatory elements such as promoters and enhancers is essential for the precise regulation of gene expression by transcription factors (TFs), and changes in chromatin accessibility mediated by Chr21-encoded chromatin remodelers BRWD1 and HMGN1 have been implicated in DS phenotypes^26,27^. Therefore, we used scMEGA^28^ to integrate our ATAC-seq (chromatin accessibility) data with the RNA-seq dataset, to predict cis-regulatory elements and TFs relevant for transcriptional deregulation in DS (Methods, Fig. 3a). We then used publicly available ChIP-seq data to validate predicted TF-cis-regulatory element interactions, and experimentally validated Protein-Protein-Interaction (PPI) data to predict interactions of TFs with Chr21 genes that may explain altered TF activities (Methods, Fig. 3a).

**Fig. 3.**
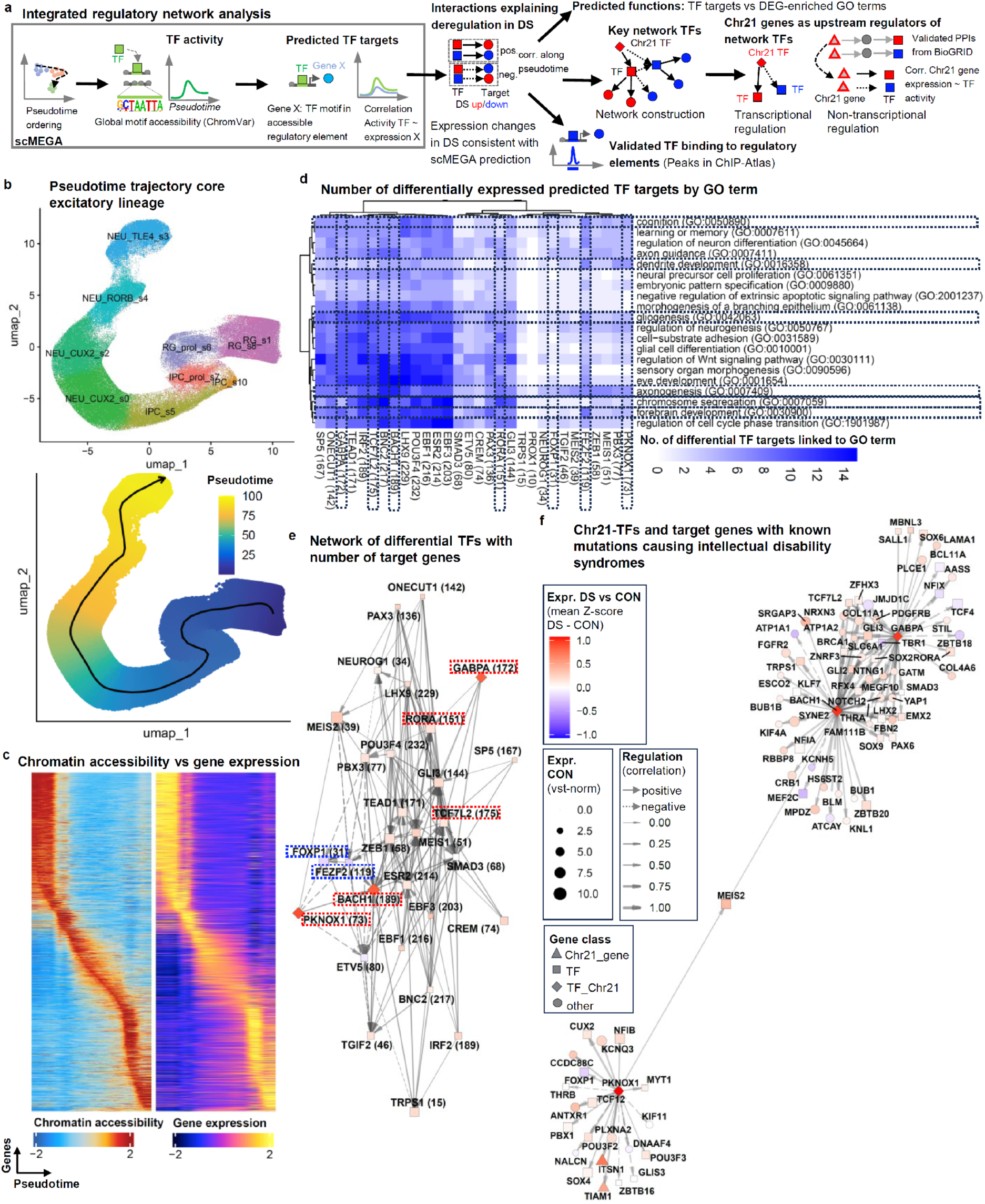
Integrated gene regulatory network analysis predicts key mediators contributing to the deregulation of transcriptional programs downstream of Chr21 genes. **a**, Approach to identify key regulators of transcriptional programs altered in DS. **b**, Excitatory lineage trajectory defined by scMEGA for network modelling. **c**, Chromatin accessibility vs gene expression along scMEGA trajectory, identifying dynamically accessible putative cis-regulatory elements indicating TF activity and determining gene expression. **d**, TFs predicted to regulate differentially expressed genes linked to altered neural functions. Heatmap showing number of interactions between TFs and differential genes linked to selected enriched gene ontology (GO) terms. In brackets: total number of targets per TF. Highlighted: GO terms and predicted key regulators discussed in main text. **e**, Network plot showing predicted interactions between TFs regulating differentially expressed genes (in brackets: number of TF-target-interactions). Node size: relative expression in control (CON) samples (vst-normalized); node color: relative expression (z-score) in DS vs CON (each mean of all cell clusters). **f**, Predicted direct targets of Chr21 TFs BACH1, PKNOX1 and GABPA with known mutations causing intellectual disability syndromes (from Genomics England PanelApp^39^). See also Extended Data Fig. 7, 8, Supplementary Table 4.

Harnessing the main excitatory populations from the whole dataset (PCW10-20), scMEGA revealed strong correlation of gene expression with chromatin accessibility at gene loci, including putative cis-regulatory elements (Fig. 3b-c). scMEGA predicted 6,299 interactions of 30 TFs with putative cis-regulatory elements of 353 differentially expressed genes with dynamic expression and accessibility along the excitatory lineage trajectory (Supplementary Table 4). Of these, 3,722 interactions were consistent with roles in determining the differences between DS and CON, i.e. for predicted positive interactions, the regulator and its target gene were both upregulated in DS or both downregulated, while for negative interactions, the regulator was upregulated in DS when the target was downregulated, or vice versa (Fig. 3a). Many of these interactions were predicted to regulate genes associated with enriched GO terms related to neural development (Fig. 3d) and included TFs that are well-characterized regulators of neuronal subtype specification and maturation, such as FEZF2, a key regulator of lower layer cortical excitatory neuron specification^29,30^, TCF7L2^31–33^, RORA, which is closely related to the cortical L4 excitatory neuron marker RORB and required for dendritic maturation of these cells^34^, and FOXP1, both enriched in the neuronal population reduced in DS (Fig. 1). Importantly, the network also included three Chr21 TFs, which have been implicated in mitochondrial function and stress responses, but whose roles in DS are not well understood: BACH1^35^, GABPA^36^, and PKNOX1^37,38^. The Chr21 TFs were predicted to directly regulate several key network nodes implicated in neural development, including FEZF2, FOXP1, RORA and TCF7L2 (Fig. 3e). Remarkably, the predicted Chr21 TF target genes were significantly enriched for genes with known mutations causing intellectual disability syndromes, as catalogued in the Genomics England PanelApp database^39^ (84 of 312 targets; odds ratio 2.0, p = 2.1×10^−7^, Fisher’s exact test), including TFs such as FOXP1, TCF7L2, SOX9 and MEF2C, and effector genes including FGFR2, NRXN3, NOTCH2 and the potassium channels KCNH5 and KCNQ3 (Fig. 3f).

As not all TFs bind to all accessible predicted binding sites, we compared our predictions with experimentally validated TF binding sites from ChIP-seq data from various human cell types in the ChIP-Atlas database^40^. This showed a significant enrichment of binding of 27 predicted upstream TFs to regulatory elements of their targets (Methods, Extended Data Fig. 7a, Supplementary Table 4), providing an experimental validation of 1419 predicted interactions (20-80% of all predicted interactions for most TFs), including several potentially important examples such as PKNOX1 targeting FEZF2, BACH1 targeting SMAD3 and SOX2, and GABPA targeting SOX9.

**Extended Data Fig. 7.**
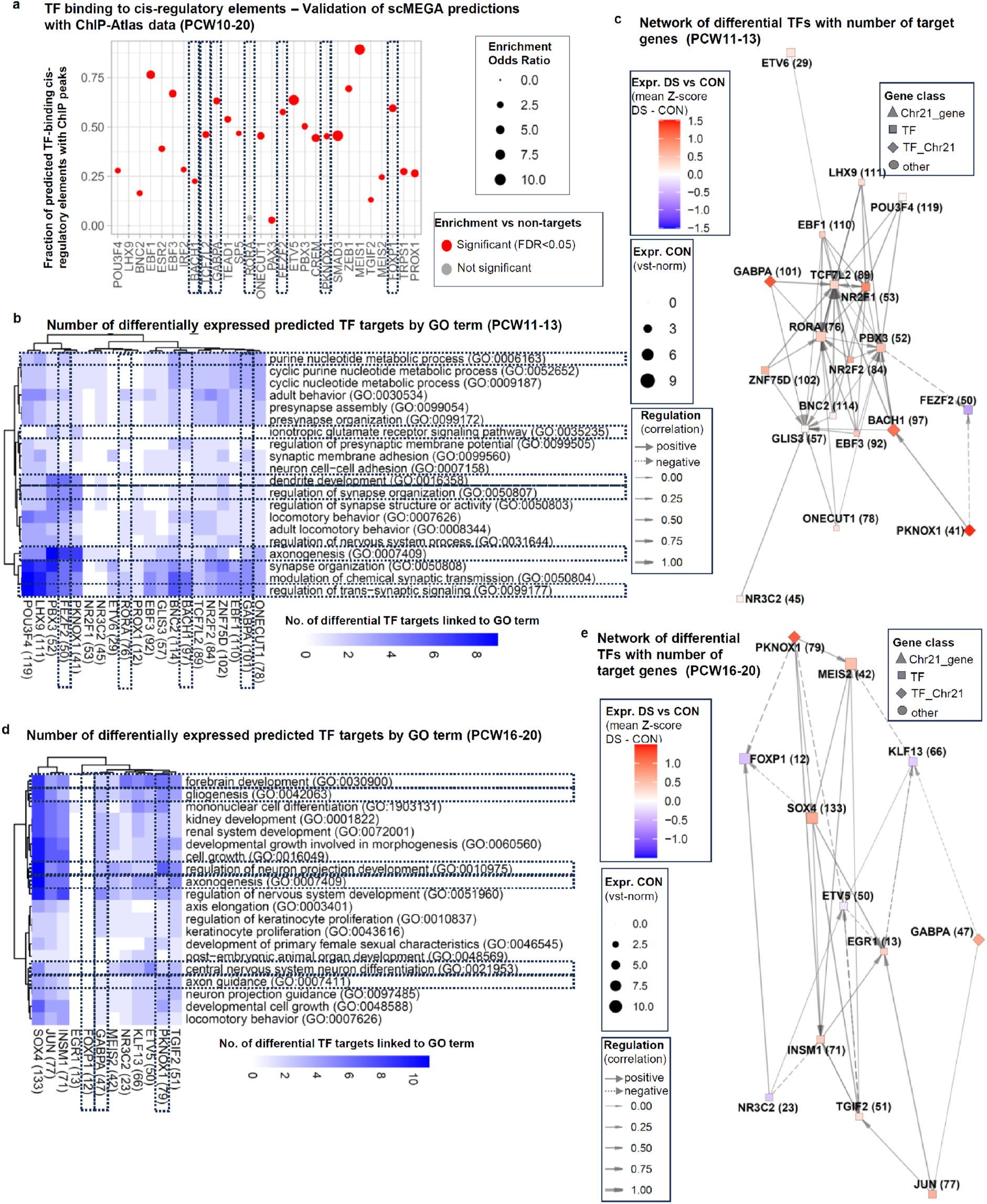
Validation of predicted TF binding to cis-regulatory elements and prediction of early and late-stage TF networks. **a**, Fraction of predicted TF-cis-regulatory-element interactions validated by published ChIP-seq data from ChIP-Atlas, with enrichment over non-targets. Note that low validation rate may be due to ChIP data being low quality or coming from non-neural cells; Boxes: selected key network TFs. **b**, TFs predicted to regulate differentially expressed genes linked to altered neural functions at PCW11-13. Heatmap showing number of interactions between TFs and differential genes linked to Top20 enriched gene ontology (GO) terms. In brackets: total number of targets per TF. Highlighted: GO terms and putative key regulators discussed in main text. **c**, Network plot showing predicted interactions of TFs regulating differentially expressed genes for PCW11-13 (in brackets: number of TF-target-interactions). Node size: relative expression in control (CON) samples (vst-normalized); node color: relative expression (z-score) in DS vs CON (each mean of all cell clusters). **d, e**, TFs predicted to regulate differentially expressed genes linked to altered neural functions, and network plot showing predicted interactions of TFs regulating differentially expressed genes at PCW16-20 (as b, c).

Separate analyses of early and late-stage samples suggest that the TF networks are highly dynamic and context dependent (Extended Data Fig. 7b-e). While PKNOX1 and GABPA emerged as key network nodes at both stages, their predicted targets differed, for instance, PKNOX1 was linked to FEZF2 in PCW11–13 samples and to FOXP1 in PCW16–20 samples. Many Chr21 genes implicated in DS phenotypes are not TFs, but the encoded proteins may modulate this TF network via PPIs. To explore this, we retrieved experimentally validated PPIs between differentially expressed Chr21 proteins and key network TFs from the BioGRID database. We then evaluated whether the expression of these interacting Chr21 proteins correlated with TF activity along the excitatory lineage trajectory identified by scMEGA (dataset PCW10-20). This would indicate that upregulation of these Chr21 genes could cause TF activity changes in DS through these PPIs (Methods, Extended Data Fig. 8, Supplementary Table 4). A total of 783 Chr21 protein-TF interaction pairs showed significant correlation, with 186 pairs, involving 35 Chr21 proteins and 22 transcription factors, showing consistence with driving changes in TF activity in DS. Notably, this approach predicted DYRK1A, one of the best-characterized mediators of DS phenotypes, as a central regulator of the TF network, influencing key network TFs including FEZF2, FOXP1, ESR2 and PAX3. APP, a Chr21 gene best known for its role in Alzheimer’s disease, but also implicated in neurodevelopment, may regulate FEZF2, GLI3, ETV5 and MEIS2, while the chromatin remodeler BRWD1 may regulate FEZF2, GLI3, ESR2 and FOXP1. USP25, a deubiquitinase recently implicated in DS-related intellectual disability^41^ may also regulate FOXP1, EBF3, BNC2 and IRF2. Only few of the identified interactions were direct (e.g. APP-FEZF2), while many were mediated by transcriptional/epigenetic co-regulators such as CREBBP(CBP) or TAF1 (Extended Data Fig. 8b, Supplementary Table 4).

**Extended Data Fig. 8.**
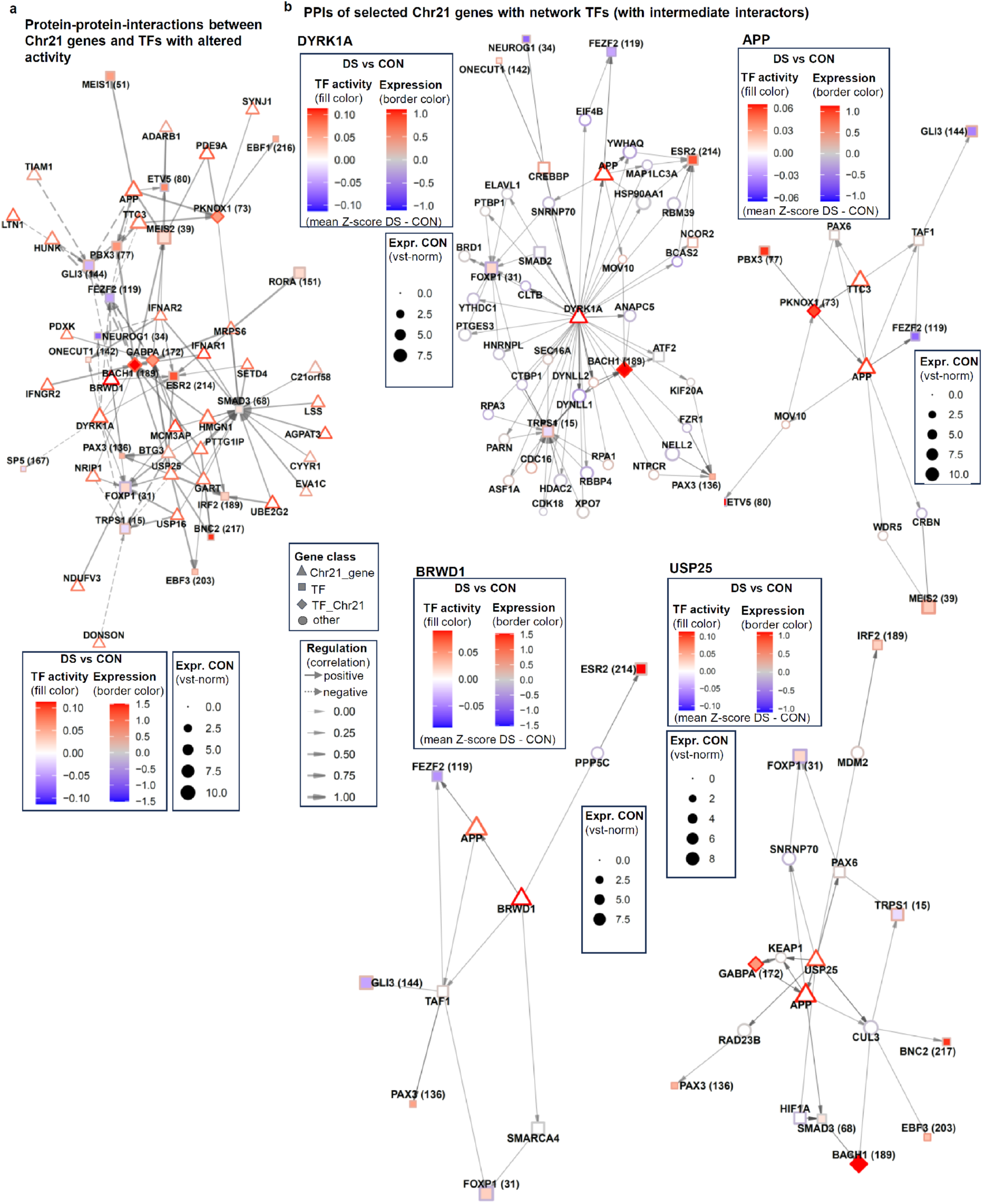
Prediction of Chr21 genes regulating network TFs via protein-protein-interactions from BioGRID database. **a**, Interactions of Chr21 proteins with network TFs predicted to regulate TF activity via experimentally validated protein-protein-interactions (PPIs) from BioGRID database. **b**, Network plot with proteins predicted to mediate interactions of Chr21 genes DYRK1A, APP, BRWD1 and USP25 with network TFs responsible for modulating TF activity changes in DS.

Together, these analyses suggest that changes in transcriptional programs in the human cortex are controlled by Chr21 TFs BACH1, PKNOX1 and GABPA, which directly regulate genes associated with intellectual disability and a network of neurodevelopmental TFs including FEZF2, FOXP1, TCF7L2, and RORA. Other Chr21 genes, such as DYRK1A, APP, BRWD1 and USP25, may also modulate these TFs via PPIs.

### Altered transcriptional programs and predicted Chr21 TF targets in the developing DS cortex are partially recapitulated in vitro and rescued by TF modulation

To further validate the mechanisms identified here, we initially assessed to what extent experimentally accessible induced pluripotent stem cell (iPSC)-based models across different differentiation stages and genetic backgrounds recapitulate human fetal cortex development and DS-associated changes.

We differentiated multiple batches of neural progenitors and neurons from two pairs of trisomic iPSC lines (named DS1, C13) from DS individuals and corresponding isogenic disomic control lines (DS2U and C9, respectively)^42–44^, and performed bulk-RNA-seq (Fig. 4a, Methods). Gene expression in cultures of *in vitro* neural progenitors (iNPCs) and neurons (iNEUs) strongly correlated with NPCs (RG, IPC populations) and neurons in the fetal cortex (Extended Data Fig. 9a), confirming successful differentiation. We detected each about 2000-4000 up- and downregulated genes in both iNPCs and iNEUs from both pairs of iPSC lines, including ∼80-100 mostly upregulated Chr21 genes (Extended Data Fig. 9b, Supplementary Table 5), indicating as expected lower variability of the side-by-side differentiated isogenic DS and CON NPCs compared to fetal tissue. Up to ∼50–90% of the differentially expressed genes (down- and upregulated) detected in fetal tissue populations were concordantly altered in NPCs *in vitro*, and up to ∼40–80% showed concordant changes in neurons, including many genes implicated in forebrain development (Fig. 4b-c). Importantly, these included also PKNOX1, BACH1, and GABPA, the Chr21 TFs predicted to be critical regulators of neurodevelopmental alterations in DS, as well as many of their putative targets (Supplementary Table 5).

**Fig. 4.**
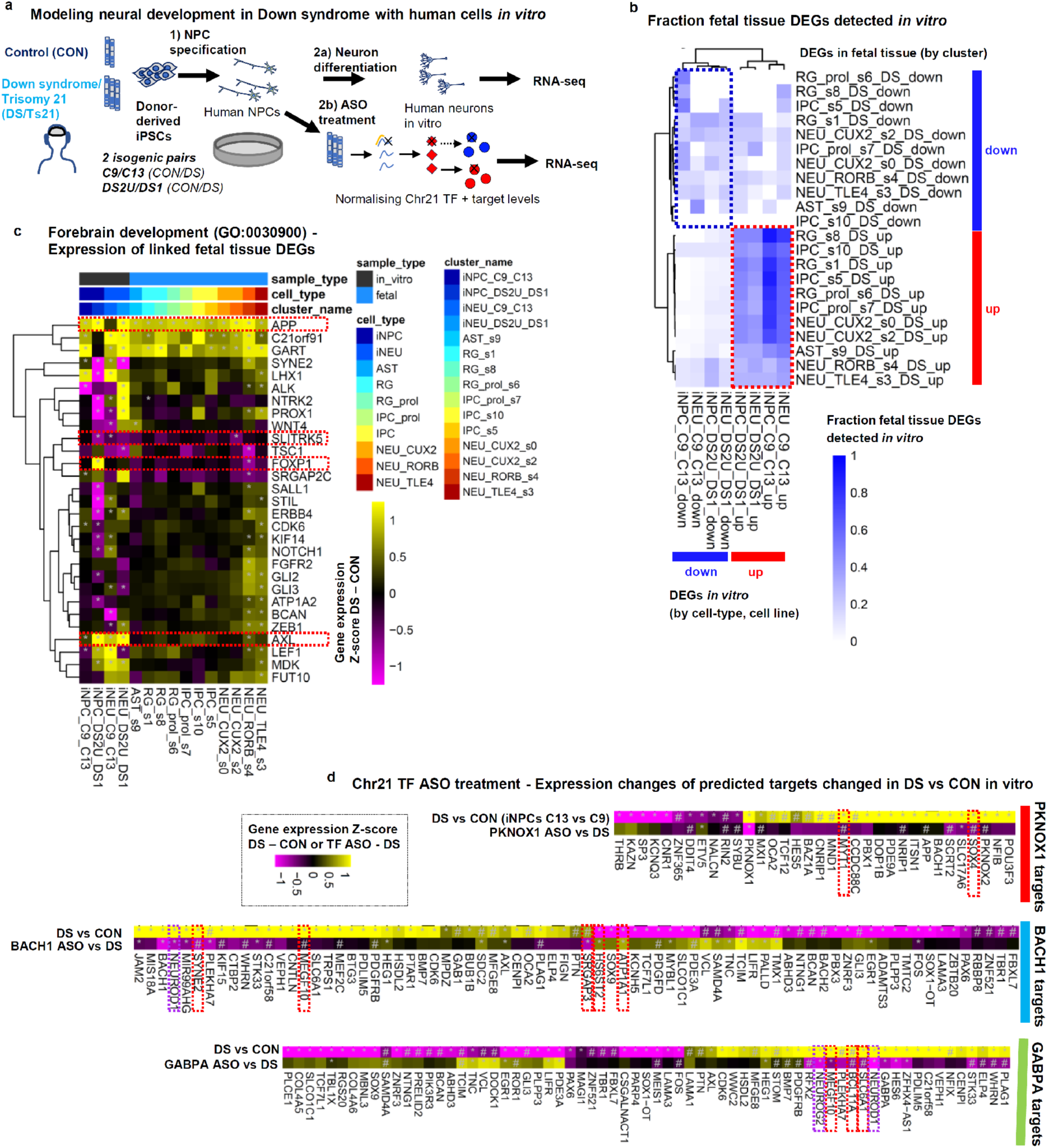
Altered transcriptional programs and predicted Chr21 TF targets in the developing DS cortex are partially recapitulated *in vitro* and rescued by TF modulation. **a,** Experimental approach for modelling DS neurodevelopment and normalizing Chr21 TF expression *in vitro*. **b,** Fraction of differential genes per tissue population also detected *in vitro* (red / blue boxes: genes up / down both in fetal tissue and *in vitro*). **c**, Expression changes in DS vs CON for genes differentially expressed in fetal tissue linked to GO term ‘*forebrain development’;* DESeq2 analysis using the Likelihood-Ratio-Test (LRT) to assess group effect across paired DS vs CON technical and biological replicates (3-10 RNA samples from wells of paired side-by-side differentiated DS/CON cells per condition from n = 6/2/3/1 independent differentiation experiments for iNPC_C9_C13/iNPC_DS2U_DS1/iNEU_C9_C13/iNEU_DS2U_DS1; see Methods and Supplementary Table 5); heatmap shows the difference in mean z-scores between DS and CON samples for each cell line and differentiation stage, and for each tissue population pseudobulk (excitatory lineage, PCW10–20); Red dotted boxes: regulators discussed in main text; *Benjamini-Hochberg-adjusted p<0.10. **d**, Expression changes of predicted Chr21 TF targets deregulated in DS-derived neural progenitors upon Chr21 TF 100nM ASO treatment; DESeq2 analysis for cultures with LRT to assess ASO effect across technical and biological replicates (7-16 RNA samples from ASO-treated and untreated wells of paired side-by-side differentiated C13 (DS) cells per condition from each n = 5 independent differentiation experiments; see Methods and Supplementary Table 5); Red/magenta dotted boxes: examples of dysregulated targets rescued by ASO treatment, including ID-linked genes (red) or other key neurodevelopmental regulators (magenta); *: Benjamini-Hochberg-adjusted across predicted TF targets p<0.10, #: trend with nominal p-value <0.1.

**Extended Data Fig. 9.**
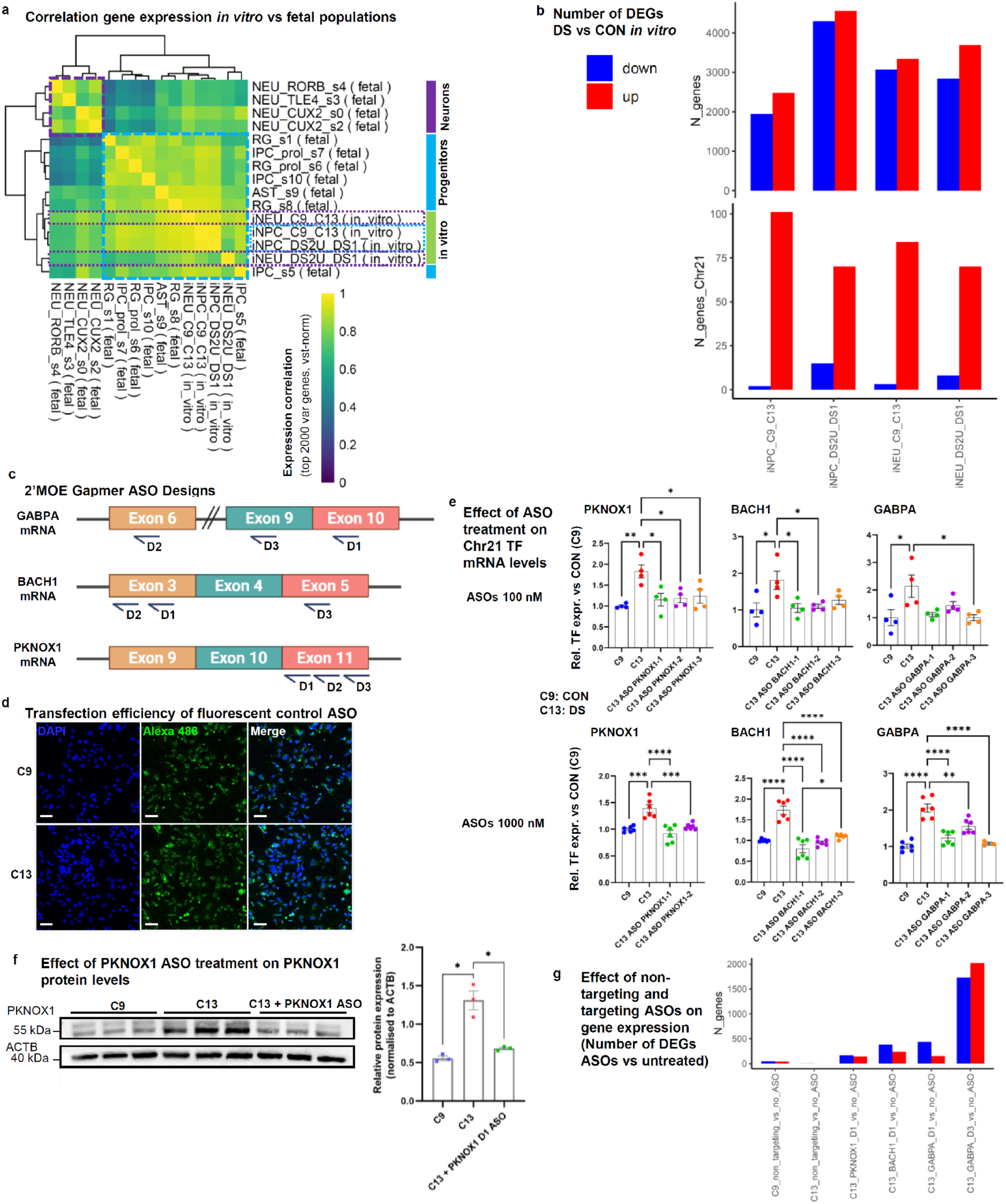
Characterization of iPSC-derived neural cells in vitro and validation of antisense oligonucleotide (ASO) mediated reduction of Chr21 TF expression. **a**, Correlation of gene expression from in vitro-differentiated neural progenitor cell (iNPC) and neuron (iNEU) cultures and in fetal tissue; Heatmap showing the correlation of mean vst-normalised expression between merged bulk data from in vitro cultures (3-10 RNA samples from wells of paired side-by-side differentiated DS/CON cells per condition from n = 6/2/3/1 independent differentiation experiments for iNPC_C9_C13/iNPC_DS2U_DS1/ iNEU_C9_C13/iNEU_DS2U_DS1; see Methods and Supplementary Table 5) and each tissue cluster pseudobulk (excitatory lineage, PCW10–20). **b**, Number of differentially expressed genes between DS vs CON *in vitro*; Benjamini-Hochberg-adjusted across predicted TF targets p<0.10, determined by DESeq2 analysis with Likelihood-Ratio-Test (LRT) to assess group effect across paired DS vs CON technical and biological replicates (3-10 RNA samples from wells of paired side-by-side differentiated DS/CON cells per condition from n = 6/2/3/1 independent differentiation experiments for iNPC_C9_C13/iNPC_DS2U_DS1/ iNEU_C9_C13/iNEU_DS2U_DS1; see Methods and Supplementary Table 5). **c,** Design of 2’-O-methoxyethyl (2’MOE) ASOs targeting Chr21 TFs. **d**, Validation of transfection of neural progenitors (NPCs) with non-targeting fluorescent ASOs (Alexa488-labelled Hs HPRT control ASO (100 nM)); Scale bar: 50 μm. **e**, qRT-PCR analysis of Chr21 transcription factor (TF) mRNA levels in hiPSC-derived NPCs from DS hiPSC line (C13) and isogenic control (C9) following treatment with 100 nM or 1000 nM of TF-targeting antisense oligonucleotides (ASOs). Expression levels were normalized to GAPDH as the reference gene. Statistical analysis was performed using one-way ANOVA followed by Tukey’s multiple comparison test (*p < 0.05, **p < 0.01, ***p < 0.001). **f**, Validation of PKNOX1 protein downregulation after 100 nM ASO treatment; Western blot with ACTB as housekeeping gene/loading control; Protein band intensities were quantified using ImageJ. Statistical analysis was performed using Welch’s t test; *\*p* <0.05, \*\**p* < 0.01, \*\*\**p* < 0.001. **g**, Non-targeting ASOs exert minimal influence on gene expression compared with chromosome 21 TF-targeting ASOs. Number of differential genes determined by DESeq2 analysis with LRT to assess ASO effect across each 2-3 technical replicates from one differentiation experiment; DESeq2 analysis with LRT test, Benjamini-Hochberg-adjusted across all genes p<0.10.

We therefore hypothesized that reducing the increased expression of these TFs in DS NPCs may rescue the dysregulated expression of their target genes and DS-associated molecular phenotypes *in vitro*. To normalize the elevated expression of Chr21 TFs and identify which predicted genes are bona fide downstream targets, we developed an antisense oligonucleotide (ASO)–based approach, a strategy successfully applied in therapies such as for spinal muscular atrophy^45^. We designed ASOs to downregulate each TF by targeting their mRNAs and established an efficient transfection protocol for iPSC-derived NPCs using fluorescently labelled non-targeting control ASOs (Extended Data Fig. 9c-d). Using quantitative reverse-transcriptase-polymerase-chain-reaction (qPCR), we confirmed effectiveness of several ASO designs at different concentrations, robustly reducing the elevated TF mRNA levels in DS NPCs close to control levels (Extended Data Fig. 9e). Western blotting for PKNOX1 demonstrated that ASO-mediated treatment also normalized TF protein levels (Extended Data Fig. 9f). Finally, we selected effective ASOs to test their ability to modulate expression of the predicted TF target genes using bulk RNA-seq. In an initial experiment, we confirmed that the treatment with non-targeting ASOs had only minor effects on global gene expression compared to targeting ASOs (Extended Data Fig. 9g). As expected from inherent differences between *in vitro* NPCs and neural cells in tissue, and the complexity of gene-regulatory network dynamics, only a subset of predicted targets showed differential expression in DS versus control NPCs, and a proportion of these exhibited partial normalization following ASO-mediated TF modulation (Fig. 4d). Importantly, the ASO treatment reverted or showed trends towards reverting the deregulation of several predicted targets linked to intellectual disability syndromes, including the PKNOX1 targets MYT1^46^, SOX4^47^ and ETV5^48^, eight BACH1 targets, including HS6ST2^49^, LIFR^50^, SYNE2^51^, SRGAP3^52^ and ATP1A1^53^, and eight GABPA targets including EGR1^54^, DOCK1^55^, BCL11A^46^, MEGF10^56^ and SLC6A1^57^, as well as previously established regulators of neuronal differentiation, such as the BACH1/GABPA target NEUROD1 and the GABPA target NEUROG2^58^.

Together, these results suggest that iPSC-derived neural cells cultured *in vitro* partially recapitulate molecular DS phenotypes in the fetal human cortex, and that Chr21 TFs PKNOX1, BACH1 and GABPA partially drive these phenotypes.

### Transplanted human neural cells reveal DS molecular and cellular phenotypes not recapitulated *in vitro* and emerging at later stages of fetal development

Our *in vitro* isogenic stem cell model captured key DS neural phenotypes, though some fetal tissue gene expression patterns differed (e.g., FOXP1 upregulated in DS1 iPSC-derived NPCs (Fig. 4c); many predicted Chr21 TF targets unchanged), highlighting both its utility for mechanistic studies and the value of fetal benchmarking. However, developing therapeutic strategies targeting the identified regulators will require models that more closely mimic the *in vivo* environment of the human brain. We therefore tested to what extent our recently established DS human xenograft system^59^, which avoids some of the drawbacks of animal models (e.g. lack of Chr21) and of *in vitro* conditions (e.g. limited neuronal maturation), recapitulates human fetal development and the observed changes in DS. As in our previous work, we transplanted iPSC-derived neural cells into adult mice to minimize their integration with host networks^59^, which could otherwise complicate interpretation.

We differentiated one of the DS iPSC lines and the corresponding isogenic control (DS1/DS2U) *in vitro* to mixed progenitor/neuron cultures, which we transplanted into the brains of adult immunodeficient mice. After 12-24 weeks of maturation *in vivo*, we extracted 9 CON and 8 DS grafts and performed single-nucleus RNA-seq (Fig. 5a, Methods, Extended Data Table 6). Pooling some grafts due to low nuclei recovery and removing low quality libraries (Methods), we retained 8 CON and 4 DS datasets with a total of 120,009 nuclei. We removed nuclei with only few transcripts mapping to the human genome (<500 transcripts/cell), presumably representing low quality nuclei and/or host mouse nuclei contaminated with ambient human RNA. We mapped the retained 98,545 high-quality high-confidence human graft nuclei to our fetal tissue dataset (see Methods), to identify the most similar fetal cells (Extended Data Fig. 10a). The vast majority of 89,844 graft cells mapped to the fetal excitatory lineage, which we subsetted and re-mapped to the fetal excitatory lineage clusters (Fig. 5b, Extended Data Fig. 10a-b). Qualitative comparison of the combined graft and fetal tissue datasets revealed that grafts contained fewer progenitors and more mature RORB- or TLE4-expressing neurons than fetal tissue, suggesting a resemblance to later developmental stages. Notably, in DS grafts, we observed an increased proportion of astrocyte-like cells (AST_s9) and a reduced proportion of proliferating progenitors (Fig. 5c), consistent with previous reports at later stages of development^2^. DS grafts showed a trend toward reduced CUX2-expressing neurons (NEU_CUX_s2) (Fig. 5c), whereas in fetal tissue the reduction occurred in the RORB/FOXP1-expressing population, possibly reflecting the difficulty of disentangling maturation-stage from subtype-related differences in cell type mapping.

**Fig. 5.**
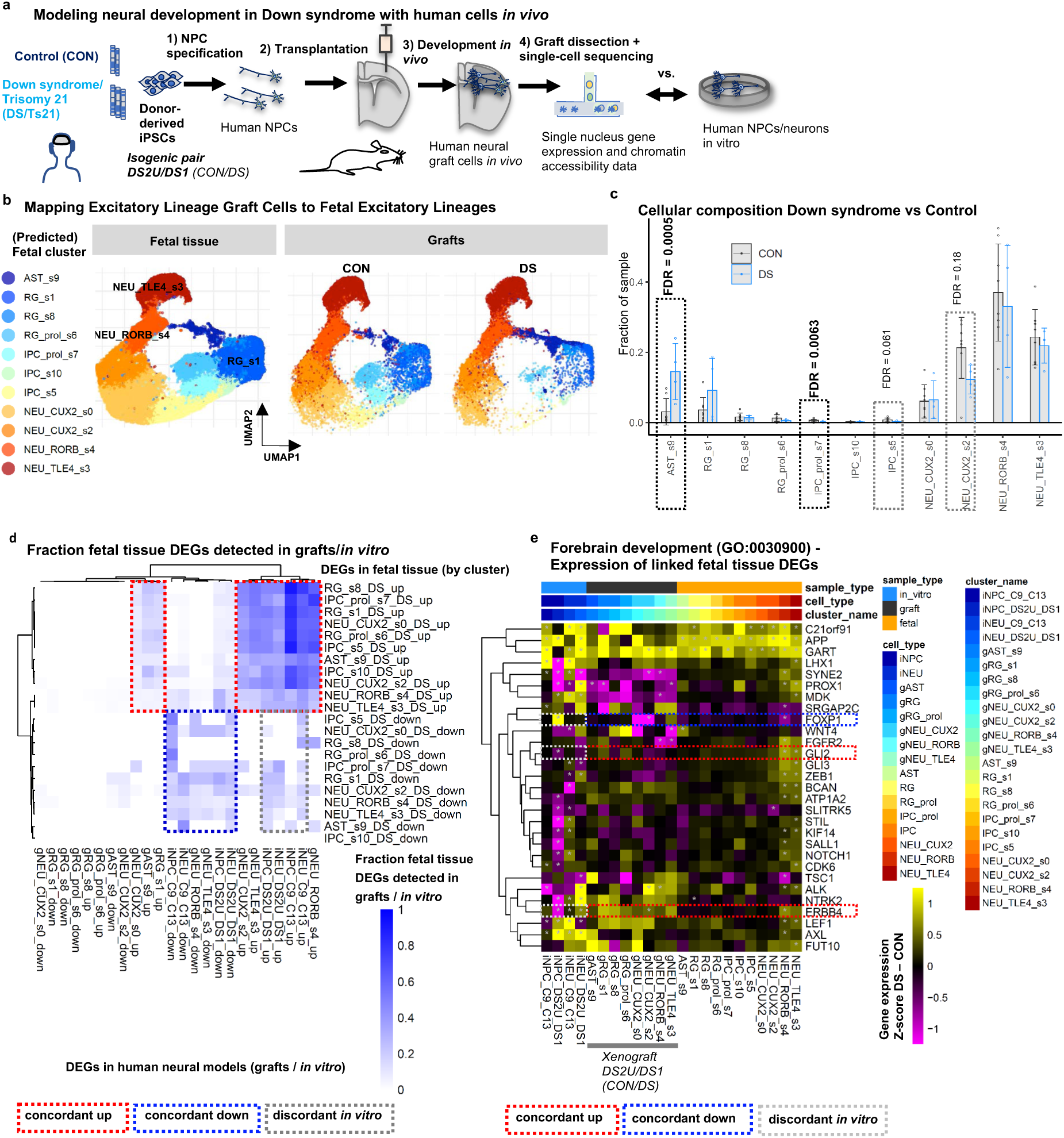
Transplanted human neural cells reveal DS molecular and cellular phenotypes not recapitulated *in vitro* and emerging at later stages of fetal development. **a,** Experimental approach for modelling DS neurodevelopment *in vivo*. **b**, Mapping of CON and DS excitatory lineage transplanted cells to fetal tissue populations (UMAP plot). **c**, Cell abundance in CON and DS transplants; Barplot showing individual samples, mean+/-sd and FDR for sccomp compositional analyses^18^ (other clusters FDR>0.05); Grey dotted boxes highlight selected populations: FDR < 0.05 (black) or FDR > 0.05 (grey). **d**, Fraction of differential genes per cluster in fetal tissue also detected in grafts and *in vitro*; highlighted: comparisons showing concordant changes in fetal tissue vs models, and increased fraction of discordantly regulated genes *in vitro* vs fetal tissue; DESeq2 analysis for grafts with LRT test by cluster with correction for sequencing technology, threshold padj < 0.10. **e**, Expression changes in DS vs CON for genes differentially expressed in fetal tissue and linked to GO term ‘*forebrain development’*; heatmap shows difference of mean z-scores between DS and CON samples (corrected for sequencing technology) for merged bulk data from in vitro cultures (3-10 RNA samples from wells of paired side-by-side differentiated DS/CON cells per condition from n = 6/2/3/1 independent differentiation experiments for iNPC_C9_C13/iNPC_DS2U_DS1/ iNEU_C9_C13/iNEU_DS2U_DS1; see Methods and Supplementary Table 5) and for each cluster pseudobulk from the graft and fetal tissue analyses (excitatory lineage, PCW10-20); Grey asterisks indicate padj < 0.10. See also Extended Data Fig. 10, Supplementary Table 6.

**Extended Data Fig. 10.**
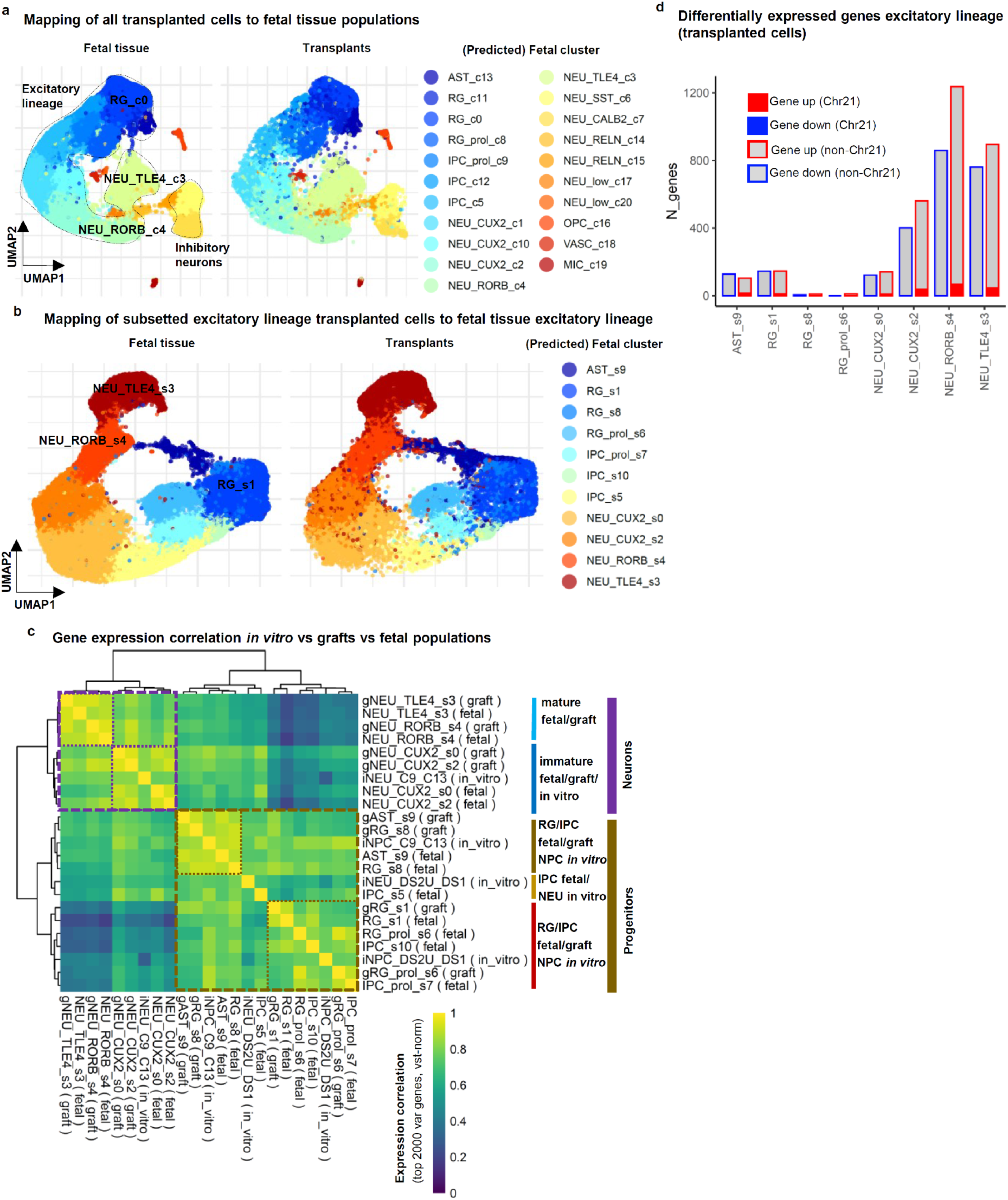
Comparison of molecular phenotypes in iPSC-derived neural cells from transplants and *in vitro* with fetal DS cortex. **a**, Mapping transcriptomes of all transplanted cells to fetal tissue populations (UMAP plot). **b**, Mapping transcriptomes of subsetted excitatory transplanted cells to PCW10-20 fetal tissue excitatory lineage populations (UMAP plot). **c**, Correlation of gene expression in neural cells *in vitro* with graft and fetal tissue populations; heatmap shows correlation of mean vst-normalised expression between each *in vitro* experiment (3-4 technical replicates) and each cluster pseudobulk from the transplants and fetal tissue analyses. **d**, Number of DEGs between DS vs CON in grafts; DESeq2 analysis for transplants with LRT test by cluster with correction for sequencing technology, threshold padj<0.10.

Importantly, expression of the most variable genes correlated more strongly between fetal and corresponding transplanted cell populations than between fetal and cells cultured *in vitro* (Extended Data Fig. 10c), suggesting as expected that transplanted cells are more similar to the fetal tissue cells than cells *in vitro*. 3290 genes were differentially expressed between DS and CON grafts, mostly in neuronal populations as in fetal tissue (Extended Data Fig. 10d). Up to >70% of genes upregulated in fetal cells were concordantly upregulated in the corresponding graft populations, while less genes were discordantly regulated compared to the *in vitro* dataset (Fig. 5d). Importantly, several genes associated with forebrain development that were dysregulated in fetal tissue showed consistent changes in transplanted cells, but discordant changes *in vitro*, including ERBB4, ALK, and FOXP1 (Fig. 5e).

Altogether, this suggests that transplanted cells recapitulate many molecular and cellular DS phenotypes - including phenotypes not recapitulated in vitro and phenotypes emerging at later stages of fetal development for which human tissue samples are scarce.

## Discussion

Our results provide insights into the mechanisms underlying alterations in fetal cortical development that may contribute to intellectual disability and cognitive impairment in DS, one of the most common congenital causes of lifelong disability. We mapped the cell type-specific transcriptional landscape of the DS fetal cortex during PCW10-20, identifying a reduction in RORB/FOXP1-expressing layer 4-like excitatory neurons. This phenotype mirrors findings in adults with DS^11^, who are predisposed to AD–like dementia. Notably, these neurons are also reduced in AD^16^, yet their early deficit in our study suggests that in DS, this reduction likely reflects impaired generation or maturation rather than neurodegeneration. We further detected changes in ∼ 700 genes within the excitatory lineage, including genes involved in forebrain development, excitatory neuron subtype specification, and genes mutated in intellectual disability. These represent some of the earliest known molecular and cellular phenotypes in the human cortex that may be associated with clinical features of DS, including intellectual disability. Interestingly, many of the observed changes seem to persist into adulthood, including the reduction of RORB-expressing neurons reported also in the adult DS cortex^11^, and a large fraction of deregulated genes also identified in a bulk-transcriptomics study of the adult cortex^12^. Our resource complements previous single-cell analyses of typical brain development^15,60–64^ by adding more samples and chromatin accessibility data from this critical period.

Remarkably, but in line with findings in the adult DS cortex^11^, we found only subtle changes in the deregulated genes. This suggests that in DS, despite the increased dosage of more than 200 genes on Chr21, not all show upregulation in cortical cells, and this does not result in pronounced deregulation of a small subset of key downstream genes that would emerge as obvious therapeutic targets. Instead, numerous subtle molecular changes appear to act synergistically, affecting shared pathways and biological processes - resembling the architecture of many idiopathic disorders, where multiple small-effect risk alleles and environmental influences collectively drive pathogenesis. In this context, given the likely limited effect of targeting single effector proteins, a more impactful therapeutic strategy could be to alter upstream transcription factors. This would allow for the coordinated regulation of numerous effector genes at once, providing a more robust method for correcting the deregulated transcriptional state^65^. To identify such master modulators, we implemented a novel multimodal network analysis approach, which revealed a regulatory network predicted to influence many of the subtle changes observed in the DS fetal cortex. This network includes 30 transcription factors predicted to interact with 6,299 accessible chromatin regions, representing putative active cis-regulatory elements of 353 deregulated genes. We identified three TFs on Chr21 that have not received much attention in the context of DS, PKNOX1, BACH1 and GABPA, as potential critical drivers of transcriptional programs altered in DS, that may contribute to the neurodevelopmental alterations resulting in intellectual disability. Their inferred targets include TFs whose disruption has been shown to impair excitatory neuron subtype specification, including FEZF2^29,30^, FOXP1^20^ and RORA^34,66^, suggesting a mechanism underlying the selective deficit in RORB/FOXP1-expressing excitatory neurons. Targets also include over 80 genes with known mutations causing intellectual disability syndromes, suggesting that their deregulation could collectively drive the DS-associated intellectual disability. Supporting our predictions, these Chr21 TFs and several of their predicted targets, including FOXP1, RORA, GLI3 and CUX2, have also been highlighted as potential key regulators of altered neurodevelopmental programs in a recent preprint using complementary analysis approaches on fetal cortical tissue^67^. As an internal validation of our approach, our analysis independently identified *DYRK1A*—a Chr21 gene already under clinical investigation for DS^68^, alongside other previously implicated candidates such as APP^69^, BRWD1^26^, and USP25^41^, which may contribute to altered neurodevelopmental programs by modulating key neurodevelopmental TFs through PPIs.

We also demonstrated that stem cell-derived human neural cells from two individuals with DS, cultured *in vitro* and compared to isogenic controls, recapitulate many molecular changes observed in the fetal brain—including alterations in PKNOX1, BACH1, GABPA, and their predicted targets. These results support our findings and underscore the value of this experimentally accessible model for studying the function of these hubs and further investigating their roles.

Using ASOs, a novel class of therapeutic agents which have delivered groundbreaking treatments for genetic disorders such as spinal muscular atrophy^45^, we were able to normalize the elevated expression of Chr21 TFs and partially revert DS-associated changes in the expression of several predicted target genes associated with neuronal differentiation and intellectual disability. Given that in many cases, including the Chr21 TF targets SOX4 and ATP1A1 validated here, even haploinsufficiency, of a single intellectual disability gene can cause cognitive impairment in humans^47,53^, our findings that the Chr21 TFs regulate many of these genes suggest a mechanistic link between chromosome 21 transcriptional dosage and the neurodevelopmental pathology of DS.

Finally, we demonstrated that our previously established xenograft model^59^, despite the heterochronic design, recapitulates key cellular and molecular phenotypes of DS, including late developmental features and *in vivo* phenotypes that extend and complement our *in vitro* benchmarking. Given the scarcity of human tissue at these late developmental stages, this model offers a valuable preclinical platform for investigating the mechanisms underlying altered cortical development in DS. While we observed a significant reduction in L4/ FOXP1+ neurons in primary DS tissue, this phenotype was not fully recapitulated in our model systems. This likely reflects the known challenges of *in vitro* models in generating upper-layer neuronal diversity and the technical complexities of mapping cell subtypes across different maturational states in xenografts. This highlights that fully modeling the nuanced process of cortical layer specification remains a key challenge for the field and an important area for future model development. Although some xenograft phenotypes could emerge after extended *in vitro* maturation, maintaining cultured neurons for 12-24 weeks, as in the xenografts, is extremely challenging. Additionally, our xenograft model provides a unique system to evaluate ASO tools targeting human candidate genes identified in this study to rescue human-specific cellular phenotypes^59,70^ *in vivo*—tools that require engagement with the human gene sequence and are not effectively assessed in conventional mouse models. Although our study delivers a valuable publicly available single-cell multiomic atlas resource— identifying cell type-specific changes, associated gene regulatory networks, a proof-of-concept ASO-mediated molecular rescue and a benchmarking of human *in vitro* and *in vivo* models—several limitations should be acknowledged. These include both technical and biological factors that may influence data interpretation and highlight areas for future investigation. Surgical terminations disrupt large-scale brain architecture, making it difficult to resolve precise cortical subregions or laminar orientation in tissue samples. This contributes to variability in cellular composition, which may obscure changes in relative cell-type abundance beyond the reduction of FOXP1/RORB-expressing neurons in single nucleus analyses. While we successfully validated this phenotype using FOXP1, including in well-preserved paraffin-embedded human brain sections, this analysis was constrained by technical limitations, as commercially available RORB antibodies proved unreliable. It will be important for future studies to confirm and extend these observations using additional markers for this neuronal subtype. Variable cell numbers/proportions also limit the sensitivity to detect differentially expressed genes, particularly in bulk RNA-seq analyses and for rare populations in single nucleus analyses. We also acknowledge that due to the technical challenges of generating xenografts, the benchmarking of our xenograft model is based on a single isogenic pair with imperfectly balanced experimental groups. Remarkably, despite these limitations, the paired comparison with the corresponding *in vitro* cultures suggests that, at least in this genetic background, the xenograft model appears to more faithfully recapitulate some DS-associated molecular changes than neural cells differentiated *in vitro*. Future work will be needed to validate these observations across additional genetic backgrounds and to establish more robust protocols for xenografting iPSC lines that are difficult to differentiate.

The regulatory networks predicted here were built on correlative predictions and TF-binding/PPI data from different cell types. These predictions will require experimental functional validation, as we have demonstrated here for targets of the Chr21 TFs PKNOX1, BACH1 and GABPA, whose dysregulation was partially rescued by ASO-mediated normalization of TF expression levels. While this approach requires further optimization and validation across additional models and assays, our proof-of-concept experiments show that targeted modulation of dosage-sensitive Chr21 TFs can normalize disease-associated gene expression.

A critical future direction will be to functionally validate the role of these key TFs by testing whether reversing their target deregulation can rescue core DS cellular phenotypes, including the reduced synchronized neural network activity we recently identified^59,70^. Such *in vivo* experiments in our humanized mouse model, while beyond the scope of the present study, are essential for confirming their contribution to DS pathology and to evaluate the potential of targeted or combinatorial modulation of these factors. Notably, a recent study demonstrated that a similar transcriptional network correction strategy can counteract impairments in an AD model^65^, supporting the feasibility of this approach. Future studies also need to define a temporal window in which targeting these TFs may be effective. Importantly, these three key TFs remain expressed throughout adulthood^11^, suggesting that targeting these TFs may still be beneficial at postnatal or adult stages.

In conclusion, this study generates a foundational molecular map of the DS cortex during a critical developmental window. This resource, combined with the *in vitro* and *in vivo* platforms we benchmarked, enables the identification and future preclinical validation of candidate regulators, paving the way for new strategies to address the neurological symptoms of DS.

## Methods

### Fetal tissue samples and ethics

Human fetal brain samples were collected from 19 fetuses with Ts21 and 20 euploid control fetuses aged PCW 10-20, all confirmed by karyotyping through the Human Developmental Biology Resource (HDBR), following pregnancy termination with maternal informed written consent. Human fresh-frozen brain tissue was provided by the Joint MRC/Wellcome Trust HDBR (Project Number 200585, supported by Joint MRC/Wellcome Trust grant #099175/Z/12/Z and MR/006237/1; http://www.hdbr.org), in compliance with ethical approval from the National Health Service (NHS) Research Health Authority (HDBR; London/Newcastle; REC approval 18/LO/0822 and 18/NE/0290) and stored at -80°C. The HDBR is overseen by the UK Human Tissue Authority (HTA) and operates in compliance with the applicable HTA Codes of Practice. Sample sizes for human fetal tissue were determined and constrained by the availability of this precious donated material. Since the study primarily relies on systematic global computational analyses rather than strong initial hypotheses, systematic blinding was not necessary. We matched sex and developmental stage between CON and DS syndrome groups, as far as sample availability allowed (see Supplementary Table 1).

Paraffin-embedded, immersion-fixed (4% PFA in PBS, pH 7.4) *post-mortem* human prenatal brain tissue (16–20 PCW; Extended Data Fig. 2g) was obtained from the Zagreb Neuroembryological Collection with ethical approval from the Internal Review Board of the Ethical Committee of the University of Zagreb School of Medicine. Procedures followed the Declaration of Helsinki (2000).

### Tissue sectioning and immunostaining

Fresh frozen brain tissue was cut on a cryostat (Leica) in 20 μm-sections for immunostaining, mounted on slides, or 80 μm-sections for nuclei extraction, collected in RNase-free low-binding tubes (LoBind, Eppendorf), which were stored at -80°C for further processing. For immunostaining to identify cortical tissue, sections were briefly thawed and dried (∼30 min), before fixation in 4% PFA in PBS for 15 min at 4°C. Sections were washed with PBST (PBS + 0.1% Triton-X100), incubated in blocking buffer (PBST + 10% normal goat serum) for 1h at room temperature (RT) and then with primary antibodies overnight in blocking buffer in a humidified chamber at 4°C. After 3 washes (PBST), sections were incubated in the dark at RT for 2h with secondary antibodies in blocking buffer, and again washed 3 times before incubating with DAPI (1 ug/ml in PBS) for 45 min. After one wash in PBS, the sections were embedded with ProLong™ Gold Antifade Mountant (Thermo Fisher P36930), and stored in the dark at 4°C. Primary antibodies used were mouse anti-SATB1/2 (Abcam, ab51502; 1:100), rabbit anti-PAX6 (Biolegend #901301; 1:200), rat anti-CTIP2 (Abcam, ab18465; 1:500), rabbit anti-FOXP1 (Abcam, ab16645; 1:200). Secondary antibodies used were anti-mouse-AlexaFluor-488, anti-rabbit-AlexaFluor-555, anti-rat-AlexaFluor-647 (all raised in goat, Thermo Fisher). Sections were imaged with a Leica SP8 confocal microscope (10X or 20X objectives), creating a single Z-plane scan of the whole tissue section using the Leica Application Suite X. Paraffin-embedded tissue (Extended Data Fig. 1g) was sectioned coronally at 10 μm using a Leica SM2000R microtome. Slides were deparaffinized in xylene (2 × 10 min), rehydrated through graded ethanol (100%, 96%, 70%), and rinsed in PBS. Sections were blocked (1% BSA, 0.5% Triton X-100 in PBS) for 2 h at room temperature. Primary antibodies (CTIP2, rat, ab18465 Abcam 1:500, FOXP1, rabbit, Abcam, ab16645, 1:200; SATB2, rabbit, Santa Cruz Biotechnology, sc-81376, 1:400) were applied overnight at 4°C. After PBS washes, secondary antibodies (anti-rabbit-AlexaFluor-488, anti-rat-AlexaFluor-555, Thermo Fisher) were incubated for 2 h at room temperature in the dark. Sections were treated with TrueBlack® to reduce autofluorescence, washed, then mounted with DAPI-containing Vectashield®. High-resolution images were acquired using the Hamamatsu NanoZoomer 2.0 RS scanner with a 40× (NA 0.75) objective at 455 nm/pixel. Fluorescence images were captured using the Hamamatsu LX2000 Lightning exciter and an Olympus FV3000 confocal microscope with a 20× (NA 0.75) objective, using FV31S-SW Fluoview software at 1024 × 1024 resolution.

### Quantification of FOXP1 immunostaining

Images of DAPI/SATB2/FOXP1/CTIP2 immunostainings were taken with a Leica SP8 (20X objective) with the same settings for all stained cryosections, creating a single Z-plane scan of the whole tissue section. To quantify nuclear FOXP1 intensity with high throughput and avoid manual counting bias, we developed an automated FIJI/R analysis pipeline (https://github.com/lattkem1/Nuc_fluor_Fiji_R_025). Regions of interest (ROI) of well-preserved cortical plate tissue were manually defined, unaware of genotype and FOXP1 staining, as regions with nuclear staining in the CTIP2 channel (6 ROI per section/sample). For each cortical plate ROI, nucleus ROIs were identified in the DAPI channel using automated threshold selection and a FIJI watershed algorithm to separate close nuclei, followed by measuring the mean intensity for each nucleus-ROI channel. FOXP1 fluorescence intensity per nucleus was further analyzed using R scripts. Fluorescence intensity per nucleus by experimental group was summarized and plotted as histograms. A threshold of 10,000 A.U. for positive/negative classification was determined based on these histograms and manual inspection of images, and nuclei from all ROIs of each sample of both groups (DS/CON) were compared using a 2-sided t-test. The group difference was confirmed to be robust to different positive/negative thresholds (not shown).

### Human iPSC culture and neural induction

Two pairs of human induced pluripotent stem cells (iPSCs) lines from individuals with DS and two corresponding isogenic iPSC lines were used. From WiCell, we acquired the trisomic DS1 line (UWWC1-DS1) and the corresponding isogenic disomic line DS2U (UWWC1-DS2U)^44^. The trisomic line C13 (DS) and isogenic control C9 (CON) were previously generated and described by Dean Nizetic, Ivan Alic and colleagues^42,43^.

iPSCs were maintained on Matrigel (Corning)-coated 6-well culture plates in mTeSR Plus medium supplemented with 0.5 *μ*M Thiazovivin (Tocris). Media changes were performed the next day with complete mTeSR Plus medium (STEMCELL Technologies) and then refreshed every other day.

For most *in vitro* experiments, adherent cultures of neural progenitor cells (NPCs) were derived from iPSCs using the Gibco protocol (Thermo Fisher Scientific, MAN0008031) and used between passages 6-10 (adapted by Dean Nizetic and colleagues, protocol “DN” in Supplementary Table 5). NPCs were expanded in Geltrex-coated 6-well culture plates prepared by diluting a 60 μL Geltrex aliquot in 6 mL cold DMEM/F12, incubated at 37°C for at least 60 minutes. Cells were thawed, centrifuged at 300 × g for 5 minutes, and resuspended in Neural Expansion Medium (NEM) with 5 *μ*M ROCK inhibitor Y27632 (STEMCELL Technologies) at a density of 1-2 million cells per well. Media changes were performed the next day with complete NEM and then changed every other day thereafter.

For most ASO experiments, including validation of ASO efficacy using qPCRs (Extended Data Fig. 9e, Supplementary Table 5), neural progenitor cells (NPCs) were differentiated from human induced pluripotent stem cells (hiPSCs) following a previously published protocol (Li et al.^71^, protocol “LI” in Supplementary Table 5). Briefly, hiPSCs were dissociated into single cells and plated at a density of 30,000 cells/cm² in neural induction medium (NIM), composed of DMEM/F12 and NeuroBasal (1:1) supplemented with 1% N2, 2% B27, 1% Penicillin-Streptomycin, 1% GlutaMax, 10 ng/ml human leukemia inhibitory factor (hLIF), and 5 μg/ml bovine serum albumin. The medium was further supplemented with 4 μM CHIR99021 (Tocris), 3 μM SB431542 (Sigma), and 0.1 μM Compound E (Millipore) for seven days. Cultures were subsequently passaged at a 1:3 ratio for five passages using Accutase, and maintained in NIM without Compound E on Matrigel-coated plates.

For xenotransplantation experiments, and the *in vitro* bulk RNAseq analysis using the DS2U and DS1 lines (see Supplementary Table 5), NPCs were generated using a neurosphere-based protocol developed by Su-Chun Zhang and colleagues^72^ (“SCZ” in Supplementary Table 5). iPSCs were transitioned to Neural Differentiation Medium (NDM), consisting of a 1:1 mixture of DMEM/F12 (Thermo Fisher Scientific) and Neurobasal medium (Thermo Fisher Scientific) supplemented with 1× GlutaMAX (Thermo Fisher Scientific), 0.5× N2 (STEMCELL Technologies), 0.5× B27 (STEMCELL Technologies), 100 *μ*M ascorbic acid (Sigma-Aldrich). On the next day, dual SMAD inhibition was initiated by supplementing NDM with 10 *μ*M SB431542 (Tocris) and 2 *μ*M DMH1 (Tocris). Cells were cultured for seven days with daily medium changes. Cells were then dissociated with Versene and transferred to low attachment flasks in NDM supplemented with 10 ng/mL basic fibroblast growth factor (bFGF, STEMCELL Technologies) and 0.5 *μ*M Thiazovivin to promote neurosphere formation. Neurospheres were maintained with medium changes every three days until day 25–29, after which they were prepared for transplantation (see below) and simultaneously plated for an additional 30 days in culture to allow comparison of the same cells *in vivo* and *in vitro* by RNA-seq.

### Cortical neuron differentiation

24-well culture plates, with or without coverslips, were coated with 300 *μ*L Poly-L-ornithine (Sigma-Aldrich) per well and incubated overnight at 37°C, followed by coating with 20 *μ*g/mL Laminin (Thermo Fisher Scientific) for at least two hours at 37°C.

For differentiation from adherent iPSC-derived NPCs, NPCs were dissociated with Accutase (Thermo Fisher Scientific) and seeded onto Poly-L-ornithine/Laminin-coated 24-well culture plates at a density of 70,000 cells per well in NEM with 5 *μ*M Y27532. Media change was performed the next day with complete BrainPhys Neuronal Medium (STEMCELL Technologies) and then changed every four days for two weeks. After which, BrainPhys Neuronal Medium is supplemented with 2 *μ*g/mL Laminin for media changes every four days until 30 days of differentiation.

For differentiation from day 25-29 neurospheres^72^ (iPSC lines DS2U, DS1, see Supplementary Table 5), neurospheres were dissociated using TrypLE (Thermo Fisher Scientific) and seeded onto Poly-L-ornithine/Laminin-coated 24-well culture plates at a density of 70,000 cells per well in Neuron Medium, composed of Neurobasal medium supplemented with 1× GlutaMAX, 1× B27 with vitamin A (STEMCELL Technologies), 10 ng/mL brain-derived neurotrophic factor (BDNF, Thermo Fisher Scientific), 10 ng/mL glial cell line-derived neurotrophic factor (GDNF, Thermo Fisher Scientific), 1 µM dibutyryl cyclic AMP (dbcAMP, STEMCELL Technologies), and 200 µM ascorbic acid. 0.1 µM Compound E (Merck Millipore) was added on the first day of plating and withdrawn during subsequent medium changes. Neuronal cultures were maintained with medium changes every three-four days in Neuron Medium for the first two weeks and partial BrainPhys medium changes until 30 days of differentiation.

### Human iPSC-derived cortical neuron transplantation and graft extraction

All mouse experimental procedures were approved by the Animal Care and Use Committee (IACUC) at Duke-NUS, Singapore (Protocol #1766), and ethics approved by the Institutional Review Board (NUS-IRB-2022-149), as well as by the Ministry of Health (MOH Ref: RR-2023/01) under the Human Biomedical Research Act guidance. Immunodeficient mice (NOD.Cg-PrkdcScid;Il2rgtm1Wjl/SzJ, JAX™ NSG) (n = 17, 4 females and 13 males) aged between 3-5 months were given 5% isoflurane mixed with oxygen as induction anesthesia. Craniotomies were performed over the right somatosensory cortex, as previously described^73^. Seven days before injection, neurospheres were dissociated into small spheres or cell clumps, and ∼2 × 10⁶ cells were seeded into an upright T25 flask with NDM plus 10 ng/ml bFGF, then transduced with 5 µl lentiviral vector expressing GFP under the human Synapsin-1 promoter. On the next day, medium was replaced with fresh bFGF-containing NDM. On the transplantation day, neurospheres were dissociated with TrypLE, washed with Cortex buffer, resuspended at 1 × 10⁵ cells/µl, and 1 µl injected with a glass needle using a micro syringe pump (UMP-3, World Precision Instruments), at the following stereotactic coordinates: anterior-posterior = -1.8 mm; medial-lateral = +2.8 mm; dorsal-ventral = -0.6 mm from bregma. A 5-mm diameter glass coverslip was placed over the craniotomy and sealed with cyanoacrylate tissue adhesive. The exposed skull was covered in dental cement and a metal plate placed on the left side of the skull, for positioning and monitoring the human cell transplant at the 2-photon microscope.

For extraction of grafts for single nuclei sequencing, mice were culled by cervical dislocation. The whole brain was extracted and immediately placed in ice-cold cortex buffer (125mM NaCl, 5mM KCl, 10mM glucose, 10mM HEPES, 2 mM CaCl2, 2 mM MgSO4). All steps from here on were conducted on ice. The hemispheres were dissected, the right-side cortex was removed from the midbrain, and the hippocampus was removed to reveal the underside of the graft. Directed by the fluorescence of the GFP-labelled human cells, a small (2-5 mm in diameter) square containing the graft was dissected using a scalpel before the mouse tissue was removed by carefully tearing along the edge of the graft using fine forceps. The extracted graft was placed in a low-adhesion Eppendorf before being flash frozen in liquid nitrogen. Samples were stored at -80°C until further processing.

### Nuclei isolation for single nucleus multiome analysis (RNA/ATAC-seq)

10-50 mg of fetal or graft tissue were processed using a protocol based on dx.doi.org/10.17504/protocols.io.bvhhn336. All steps were performed on ice or at 4°C with pre-chilled RNase-free buffers and tools, and up to 4 samples were processed in parallel.

After removal from dry ice, tissue was immediately suspended in Homogenization Buffer (10 mM Tris-HCl pH7.4, 320 mM Sucrose, 3 mM CaCl2, 3 mM MgCl2), supplemented freshly with 0.1% NP-40, 1 mM DTT and 1 U/µl RNAse inhibitor (Protector (Sigma #3335402001), RiboLock (Thermo Fisher, PN-EO0382), or RNaseOUT (Thermo Fisher, 10777019) RNase inhibitor, see Supplementary Table 1). The tissue was then immediately homogenized using a 1-ml dounce homogenizer and exactly after 5 min diluted with 1 volume of Homogenisation Buffer (without NP-40), filtered (30 µm mesh size) to remove large debris, and centrifuged (500g, 5 min, 4°C) to collect raw nuclei. Raw nuclei were resuspended in Homogenization Buffer (without NP-40) and mixed with 1 volume 50% iodixanol buffer (10 mM Tris-HCl pH7.4, 3 mM CaCl2, 3 mM MgCl2, 1 mM DTT, 0.5 U/µl RNAse inhibitor). This suspension was carefully overlaid on a 29% iodixanol buffer (as above + 160 mM sucrose), and centrifuged (6000g, 30 min, 4°C). The nuclei pellet was resuspended in a Lysis Buffer (10 mM Tris-HCl pH7.4, 10 mM NaCl, 3 mM MgCl2, 1% BSA, 1 mM DTT, 0.5 U/µl RNAse inhibitor, 0.1% Tween-20, 0.1% NP-40, 0.001% digitonin), and after exactly 2 min nuclei were diluted in 10 volumes wash buffer (lysis buffer without NP-40 and digitonin), and centrifuged (500g, 5 min, 4°C) to collect the nuclei. The nuclei were then resuspended in 1x Nuclei Buffer (from the “Chromium Next GEM Single Cell Multiome ATAC + Gene Expression Reagent Bundle”; 10X Genomics PN-1000283/PN-1000285; supplemented with 1 mM DTT, 0.5 U/µl RNAse inhibitor).

### Multiome library preparation and sequencing

snRNA-seq and snATAC-seq libraries were prepared from isolated nuclei by the NIHR Imperial BRC Genomics Facility using a 10X Genomics Chromium X and the “Chromium Next GEM Single Cell Multiome ATAC + Gene Expression Reagent Bundle” (10X Genomics PN-1000283/PN-1000285) according to manufacturer’s instructions. Libraries were sequenced using Illumina Nextseq2000 or Novaseq6000 sequencers.

### Basic processing and quality control of Multiome data from fetal tissue

Raw demultiplexed sequencing data (fastq files) were mapped to the human genome (GRCh38) and quantified using cellranger-arc (v2.0.2; 10X Genomics) and loaded into an R environment (R v4.3.3), using the Seurat single cell analysis package v5.1.0^74,75^ , with the Signac extension (v1.13.0) for analyzing single nucleus ATAC data^76^. To retain only high-quality datasets and cells, low quality nuclei and potential nuclei clumps were removed, i.e. nuclei with low or extremely high transcript counts (<500 or > 30,000 UMIs/cell), high counts of mitochondrial genes (>2%), low or extremely high numbers of mapped chromatin fragments (<100 or 25,000 ATAC counts/cell), or poor or unspecific chromatin fragmentation (nucleosome_signal > 2 or transcriptional start site enrichment < 1.1). Datasets with a high fraction of low-quality cells (>50%) or less than 500 retained cells were considered as low-quality datasets and completely removed. In cases of tissue samples with adequate tissue quality, that resulted in low quality datasets, Multiome sequencing was repeated with a second tissue aliquot. Overview of all removed and retained datasets, see Supplementary Table 1.

### Data integration, dimensionality reduction and mapping to Bhaduri et al., 2021 reference atlas

To account for technical variability such as sequencing depth and batch effects, RNA counts per cell were normalised using the function (with parameters) SCTransform(ncells = 3000, variable.features.n = 2000, conserve.memory = TRUE). Principal components of the normalised RNA counts were calculated with RunPCA() and used to integrate the individual sample datasets using IntegrateLayers(method = HarmonyIntegration, assay = “SCT”, orig.reduction = “pca”). As dimensionality reduction for visualization of cell populations based on transcriptome similarity, Uniform Manifold Projection (UMAP) was performed using RunUMAP(reduction = “harmony”, dims = 1:30, return.model = TRUE) and transcriptome similarity neighborhoods were detected using FindNeighbors(reduction = “harmony”, dims = 1:30).

The dataset was then mapped to the reference atlas from Bhaduri et al., 2021^15^. The processed count matrix and cell level metadata for the complete dataset from Bhaduri et al., 2021^15^, were downloaded from https://cells.ucsc.edu/dev-brain-regions/wholebrain/ (files meta.tsv, exprMatrix.tsv.gz; accessed 17/11/2023) and imported into Seurat. Cells with <750 UMI or >10% mitochondrial reads were removed, and the dataset was split by samples and processed as described above. Transfer anchors were generated using FindTransferAnchors(reference = seur_ref, query = seur, dims = 1:30, reference.reduction = “pca”), followed by mapping and label transfer with MapQuery(anchorset = anchors, reference = seur_ref, query = seur, refdata = list(cell_cluster = “cell_cluster”, cell_type = “cell_type”, area = “area”), reference.reduction = “pca”, reduction.model = “umap”).

Predicted cluster/cell type and area assignment were projected on UMAP dimensionality reductions (dataset randomly subsampled to 100,000 cells for plotting to avoid excessive plot sizes). The fraction of cells of each sample mapping to each of the brain areas of the reference dataset was plotted as heatmap, which identified three samples with high mapping to non-cortical regions (Extended Data Fig. 1e), which were removed from all following analyses.

The remaining samples were re-integrated, followed by dimensionality reduction and neighborhood-detection as above.

### Identification of cell populations and differential abundance testing in fetal multiome dataset

To identify transcriptionally similar cell populations at different resolutions, clustering was performed FindClusters(algorithm = 1) with different resolution parameter values (range 0.3 – 1.5). Cluster assignment at different resolutions, sample metadata, and expression of selected cell type markers were projected on UMAP dimensionality reductions (dataset randomly subsampled to 100,000 or 10,000 cells for plotting to avoid excessive plot sizes). Based on the mapping to the reference atlas and a curated set of cell type markers, cell clusters at a final resolution of 0.5 were assigned to cell types and labelled accordingly (Fig. 1c-d, Extended Data Fig. 2a).

Changes in cellular composition were assessed with the sccomp package (v1.7.15)^18^, using the functions sccomp_estimate(formula_composition = ∼group, .sample = sample, .cell_group = cluster_name, bimodal_mean_variability_association = TRUE, cores = 8) and sccomp_test(). As alternative approach for cluster-free differential abundance analysis, the MiloR package (v1.10.0)^17^ was used. For this, the Seurat object was converted to a SingleCellExperiment object (as.SingleCellExperiment()), followed by neighborhood detection with buildGraph(k = 50, d = 30) and makeNhoods(prop = 0.05, k = 50, d=30, refined = TRUE) and neighborhood abundance quantification and testing with countCells(samples=“sample”), calcNhoodDistance(d=30) and testNhoods(design = ∼ group). For each cell, membership in any differential neighborhood was identified, and the maximum |log2FC| of any associated neighborhood mapped onto the UMAP dimensionality reduction plot.

For analyses of the excitatory lineage, the Seurat object of the full dataset was subsetted to include only cells of samples from the relevant stages, and cell clusters of astrocytes (AST), progenitors (RG, IPC) and excitatory lineage neurons (NEU_CUX2, NEU_RORB and NEU_TLE4). This subsetted dataset was re-integrated and re-clustered as above (final clustering with resolution = 0.3 for whole dataset PCW10-20 and PCW11-13 and 0.5 for PCW16-20).

Throughout the analyses, as transcription factors we defined the 1672 genes from a curated list from the Fantom5 consortium (https://fantom.gsc.riken.jp/5/sstar/ Browse_Transcription_Factors_hg19, retrieved 21/12/2021). As Chr21 genes we defined the 210 protein-coding genes annotated with their HUGO Gene Nomenclature Committee (HGNC) symbol in the Ensembl database (Homo_sapiens.GRCh38.105.chromosome.21_220405.gff3, accessed 05/04/2022).

### Differential gene expression analysis

For differential gene expression analysis, for all cells of each sample and cell cluster, transcript (UMI) counts for each gene were aggregated into pseudobulk samples. To avoid spurious results due to low numbers of cells/UMI counts, pseudobulks with less than 10 cells, and very low expressed genes with less than average 0.1 UMI count per cell in all clusters were removed from further analysis. Subsequently, differential gene expression analysis was performed for cell clusters with at least 2 pseudobulks per condition (CON/DS) using DESeq2 v1.42.1^77^, comparing DS vs CON pseudobulks for each cluster (Wald test, design ∼cluster_group). Genes with padj(FDR) < 0.10 and |log2FoldChange| > log2(1.2) were considered as differentially expressed.

Overrepresentation analyses for gene ontology terms for biological processes, and for gene sets of the Molecular Signatures Database, was performed on the union of all differentially expressed genes, using the R clusterProfiler package v4.10.1^78^ with annotations from the DOSE(v 3.28.2) and org.Hs.eg.db (v3.18.0) packages. Enriched genes per term/gene set were overlapped with the differentially expressed genes per cluster to identify which genes related to the respective gene set were deregulated in which cluster. For heatmap representations, the package pheatmap (v1.0.12) was used. Gene z-scores were calculated over all analyzed pseudobulks, and for each cluster the mean of the z-scores of all controls and all DS samples was calculated, as well as the difference in mean z-scores (DS – CON) as measure to visualize the magnitude of the expression difference between both groups. As alternative single-cell-based differential gene expression analysis approach, the Nebula package was used^21^ (v1.5.3). The Seurat object was subsetted for each cluster to retain only cluster cells, converted to a Nebula object, using scToNeb(assay = “RNA”, id = “sample”, pred = c(“group”), offset=“nCount_RNA”), a model matrix generated using model.matrix(∼group, data=seuratdata$pred), and the differential expression results calculated using nebula(seuratdata$count, seuratdata$id, pred=df, offset=seuratdata$offset, ncore = 16).

### Chromatin accessibility mapping and gene regulatory network analysis

For accurate identification of accessible regions, peaks of ATAC reads were called for each cluster using the Signac function CallPeaks(group.by = “cluster_name”), using annotation packages BSgenome.Hsapiens.UCSC.hg38 (v1.4.5) and EnsDb.Hsapiens.v86 (v2.99.0), and refined by removing non-standard chromosome annotations (keepStandardChromosomes(pruning.mode = “coarse”)) and “blacklisted” regions that are generally excluded from ATAC-seq analyses (subsetByOverlaps(ranges = blacklist_hg38_unified, invert = TRUE)).

Peaks were then classified with Ensembl annotations using the EnsDb.Hsapiens.v86 package into peaks overlapping with exons, introns, promoter regions and other peaks (intergenic), using the functions intronicParts(), exonicParts() and promoters() to retrieve Ensembl annotations and the findOverlaps() function to identify overlapping ATAC peaks. To link peaks as putative active cis-regulatory elements to likely target genes, peaks were mapped to the closest gene promoter, using the distanceToNearest() function.

Gene regulatory network analysis was performed using the R package scMEGA v1.0.2^28^, based on the scMEGA Github analysis workflow for 10X Multiome data. Cells of the main excitatory lineage populations (excluding AST and NEU_low populations) were ordered along the excitatory lineage trajectory using the manually ordered subset clusters with AddTrajectory(trajectory = cluster_names, group.by = “cluster_name”, reduction = “umap”, dims = 1:2, use.all = TRUE).

All steps of scMEGA were based on the SCT-normalized RNA data and the ATAC peak data mapped by cluster, and the ChromVar TF activity data calculated from these (parameters tf.assay = “chromvar”, rna.assay = “SCT”, atac.assay = “peaks_by_cluster”). TF motifs from the JASPAR database (JASPAR2024 package, v0.99.6) were retrieved using getMatrixSet(x = JASPAR2024@db, opts = list(collection = “CORE”, tax_group = ‘vertebrates’, all_versions = FALSE)) and mapped to the ATAC peaks, using AddMotifs(genome = BSgenome.Hsapiens.UCSC.hg38). TF activity was calculated using RunChromVAR(). Selection of TFs was not restricted for the initial network analysis (except ecluding TFs with activity or expression 0 over the whole trajectory), to prevent exclusion of TFs with repressive activity and without prominent alterations along the differentiation trajectory. Peak-gene-links were identified with SelectGenes(), which also generated the matched chromatin accessibility - gene expression heatmap shown in Fig. 3c. The TF activity-gene expression correlation was calculated with GetTFGeneCorrelation(), limited to genes differentially expressed in DS. The predicted TF-target interactions were extracted with GetGRN(), and interactions between differentially expressed TFs and target genes with fdr ≤ 0.05 and abs(correlation) ≥ 0.3 were used for the construction of the “unfiltered” network based on correlation of chromatin accessibility/TF activity with gene expression along the scMEGA-defined excitatory lineage trajectory.

We then filtered for interactions that are consistent with the hypothesis that the change of expression of the TF determines the change of target gene expression in DS vs CON. For this, we calculated a scaled relative expression in DS vs CON for each gene over all cells (difference mean SCT-normalized expression Z-score DS cells vs CON cells, i.e. all genes with mean Z-score >0 are increased in DS), retaining only interactions with Z(TF) * Z(target) * correlation >0 (i.e. for a positive regulation/correlation along the trajectory, both TF and target are either upregulated (Z(TF) >0, Z(target) >0) or downregulated(Z(TF) <0, Z(target) <0) in DS, for negative regulation, either the TF is upregulated and the target downregulated (Z(TF) >0, Z(target) <0), or vice versa).

The number of transcription factor-target interactions per factor was quantified and the genes in the network were classified as transcription factors (“TF”), chromosome 21 genes (“Chr21”), chromosome 21 transcription factors (“TF_Chr21”). As (coarse) measure for the average relative gene expression of each gene, the mean of the vst-normalized expression (DEseq2 vst() function) of all CON pseudobulks was calculated. Network plots were generated using the ggraph package (v2.2.1), representing genes as nodes with node sizes corresponding to the mean CON gene expression (vst normalized) and the color indicating the relative expression in DS vs CON (difference of mean expression Z-score of DS samples and CON samples).

To validate scMEGA predictions of TF binding to putative cis-regulatory elements with ChIP-seq data publicly available in the ChIP-Atlas database, we downloaded merged bed files containing ChIP-seq peaks from all human cell datasets (files “https://chip-atlas.dbcls.jp/data/hg38/assembled/Oth.ALL.05.[TF].AllCell.bed”). Overlaps of ChIP-Atlas ChIP peaks with the ATAC peaks of the DS dataset were identified using findOverlaps(). ATAC peaks with ChIP-validated TF binding were overlapped with peaks predicted by scMEGA to bind the corresponding TF, to identify high confidence TF binding regulatory elements, and their target genes predicted by scMEGA. Enrichment of TF binding in predicted target genes was statistically assessed by quantifying the number of predicted targets with/without high confidence TF binding regulatory elements vs the background of predicted non-targets, followed by a two-sided Fisher’s Exact Test for each TF and Benjamini-Hochberg correction for multiple testing.

To identify putative protein-protein-interactions including the network transcription factors from the gene regulatory network analysis with Chr21 genes, we extracted experimentally validated protein-protein-interactions from the BioGRID database (v4.4.233), using the BioGRID API via the R packages jsonlite (v1.8.8) and httr (v1.4.7). We extracted all interactions including network transcription factors or differentially expressed Chr21 genes using https://webservice.thebiogrid.org/interactions/, then filtering for interactions of Chr21 genes with network transcription factors directly or via one common interacting protein.

Inspired by the scMEGA approach, for this space of potential Chr21 gene – TF interactions, we then calculated the correlation of TF activity along the excitatory lineage trajectory with the expression of the linked Chr21 gene. For this we assigned cells to 100 trajectory bins as determined by scMEGA and calculated the mean expression/TF activity per bin. The correlation for each Chr21-TF pair over all bins was determined with the R cor.test() function, and p-values adjusted by Benjamini-Hochberg multiple testing correction to identify statistically significant correlations. To identify interactions of Chr21 genes with TFs that might determine the changes of TF activity in DS, we filtered the interactions as described for the scMEGA analysis. We calculated mean Z-scores of Chr21 expression and TF activity of all DS – all CON cells for all Chr21 genes and TFs in the analysis, respectively, and retained only interactions with padj≤0.05, abs(correlation)≥0.2, abs(Z(TF activity))≥0.01, abs(Z(Chr21 gene expression))≥0.1, and consistent directions of the predicted regulation and changes in DS (Z (TF activity) * Z(Chr21 gene expression) * correlation >0, see scMEGA analysis).

Network plots were generated using the ggraph package (v2.2.1), representing genes as nodes with node sizes corresponding to the CON gene expression (vst normalised, see scMEGA analysis), border color indicating the relative expression in DS vs CON (difference of mean expression z-scores of DS samples and CON samples), and fill color the relative TF activity in DS vs CON (difference of mean activity z-scores of DS samples and CON samples).

### Bulk-RNA-seq and analyses

For bulk-RNA-seq of fetal tissue samples (see Extended Data Table 3), total RNA was extracted using the RNeasy Plus Micro kit (Qiagen) according to manufacturer’s instructions. RNA-seq was then performed by the NIHR Imperial BRC Genomics Facility using the NEBNext rRNA Depletion kit v2 (Human/Mouse/Rat) and NEBNext® Ultra™ II Directional RNA Library Prep Kit for Illumina. Libraries were sequenced in PE75 mode.

For *in vitro* differentiated neuronal cultures, cells were allowed to mature for 30 days before being dissociated with Accutase, pelleted, and flash frozen. Samples were stored in -80°C until further processing. Library preparation and bulk-RNA-seq were performed using AccuraCode® RNAseq Kit (Singleron Biotechnologies).

For ASO experiments, total RNA was extracted from NPCs using FastPure Cell/Tissue Total RNA Isolation Kit (Vazyme) according to manufacturer’s protocol. RNA samples were stored in -80°C before processing by DNBSEQ eukaryotic strand-specific transcriptome resequencing (BGI Genomics).

Gene level count matrices were generated by mapping to the human genome (GRCh38) using the pipelines nf-core/rnaseq (v.3.18.0; doi:10.5281/zenodo.1400710) or AccuraCode (v1.2.0; for Singleron data).

Differential gene expression analyses for the tissue bulk analysis was performed using DESeq2 comparing DS and CON using Wald’s test.

For analyses of *in vitro* experiments, groups were compared using DESeq2 with a likelihood ratio test (LRT), using a multifactorial design comparing group effects (DS vs CON or DS ASO-treated vs untreated) between one or more paired technical replicates across experimental batches (biological replicates). The analysis was performed using the commands DESeqDataSetFromMatrix(counts_comp, colData = meta_comp, design = ∼ batch + group) and DESeq(test = “LRT”, reduced = ∼ batch), applying a significance cutoff of padj < 0.1. For the ASO experiments, samples treated with different ASO designs targeting the same Chr21 TF were considered as technical replicates.

For visualizing expression Z-scores, the count matrices were normalized using vst(), followed by batch correction with removeBatchEffect(batch = meta$batch”) from the limma package^79^ (v3.58.1). For each group comparison, the difference of the mean of the z-scores by group were visualized as measure of the magnitude of the expression difference between both groups.

### Antisense oligonucleotide (ASO) treatment

NPCs derived from DS (C13) and isogenic control (C9) iPSC lines were seeded on Geltrex (Gibco)-coated 24-well culture plates, with or without coverslips, at a density of 70,000 cells per well in Neural Expansion Medium (NEM).

Upon reaching approximately 70% confluency, cells were transfected with 100 nM 2’-O-methoxyethyl (2’MOE) gapmer antisense oligonucleotides (ASOs) targeting Chr21 transcription factors (Integrated DNA Technologies), using Lipofectamine™ 3000 (Thermo Fisher Scientific) according to the manufacturer’s protocol. Transfection was carried out at 37°C for 96 hours, after which cells were harvested for downstream applications. ASO sequences are listed in Supplementary Table 5.

To assess and optimize transfection efficiency, C13 NPCs were seeded on Poly-L-ornithine-coated coverslips in NEM. The following day, the medium was replaced with BrainPhys Neuronal Medium, and cells were transfected with 100 nM Alexa Fluor 488-labelled HPRT control ASO (Integrated DNA Technologies) using Lipofectamine™ 3000 as described above. Following treatment, cells were fixed in 4% paraformaldehyde (PFA) in 1× phosphate-buffered saline (PBS) for 15 minutes at room temperature, washed thrice with 1× PBS for 10 minutes each, and permeabilised with 0.1% Triton X-100 in 1× PBS. Cells were then counterstained with DAPI (1:1000, Thermo Fisher Scientific), followed by 1× PBS washes. Coverslips were mounted using ProLong™ Glass Antifade Mountant (Thermo Fisher Scientific), stored in the dark at 4°C overnight, and then imaged using a LSM980 confocal fluorescence microscope (Zeiss).

To assess knockdown efficacy of ASOs by qPCR and RNA-seq experiments (Extended Data Fig. 9e, Supplementary Table 5), NPCs derived from DS (C13) and isogenic control (C9) iPSC lines were seeded on matrigel-coated plates at a density of 100,000 cells/cm^2^ in NIM without Penicillin/Streptomycin. ASOs were transfected using 100 nM or 1000 nM 2’-O-methoxyethyl (2’MOE) gapmer antisense oligonucleotides (ASOs) using Lipofectamine Stem (Thermo) according to the manufacturer’s protocol. Briefly, Lipofectamine Stem was mixed with Opti-MEM and incubated for 5 minutes, followed by a 15-minute incubation after addition of ASOs. The transfection mixture was added drop-wise to freshly seeded cells, and the medium was replaced the following day. Cells were harvested 4 days post-transfection for downstream analyses.

### Quantitative reverse transcription PCR (RT-qPCR)

Total RNA was extracted from NPCs using the FastPure Cell/Tissue Total RNA Isolation Kit, following the manufacturer’s instructions. RNA concentration and purity were assessed using a Nanodrop™ N2000 spectrophotometer (Thermo Fisher Scientific). For each sample, 500 ng of total RNA was reverse transcribed into complementary DNA (cDNA) using SuperScript™ IV VILO™ Master Mix (Thermo Fisher Scientific).

Quantitative PCR was performed using 10 *μ*L reactions containing 5 ng of cDNA template, 0.5 *μ*L of 10 *μ*M forward and reverse primers (Integrated DNA Technologies), and 5.5 *μ*L SupRealQ Ultra Hunter SYBR qPCR Master Mix (Vazyme). Reactions were run on a QuantStudio™ 5 Real-Time PCR System (Thermo Fisher Scientific) using the following cycling conditions: 95°C for 30 seconds, followed by 40 cycles of 95 °C for 1 second and 60 °C for 20 seconds. Melt curve analysis was performed to confirm amplification specificity. Relative gene expression was calculated using the ΔΔCt method, normalized to GAPDH as the housekeeping gene. All reactions were performed in technical triplicates. Primer sequences are provided in Supplementary Table 5.

### Western blot

NPCs were lysed in RIPA buffer (Thermo Fisher Scientific) supplemented with protease and phosphatase inhibitors on ice for 30 minutes. Lysates were centrifuged at 15,000×g for 15 minutes at 4°C, and the supernatant was collected for protein quantification using the BCA Protein Quantification Kit (Vazyme) following manufacturer’s protocol.

Equal amounts of protein (50 *μ*g) were mixed with 1x NuPAGE LDS sample buffer (Thermo Fisher Scientific) and 1x NuPAGE Sample Reducing Agent (Thermo Fisher Scientific), then denatured at 70°C for 10 minutes. Samples were resolved by SDS-PAGE on a 4-20% Mini-PROTEAN® TGX™ Precast Protein Gels (Bio-Rad) in 1x Tris Glycine-SDS running buffer at 100 V for 1 hour. A 250 kDa Plus Prestained Protein Marker (Vazyme) was used as the molecular weight reference. Proteins were transferred to nitrocellulose membranes (Bio-Rad) using a wet transfer system at 120 V for 1.5 hours on ice.

Membranes were blocked in 5% non-fat milk in TBST (0.1% Tween-20 in TBS) for 1 hour at room temperature on shaker, followed by overnight incubation at 4°C with primary antibodies diluted in blocking buffer. After three washes with TBST of 5 minutes each at room temperature, membranes were incubated with appropriate HRP-conjugated secondary antibodies for 1 hour at room temperature on shaker. After three additional washes, signal was visualised using SuperPico ECL Chemiluminescence Kit (Vazyme) and imaged on the ChemiDoc™ MP Imaging System (Bio-Rad). Band intensities were quantified using ImageJ and normalised to ACTB as the loading control. Antibodies used were mouse anti-Beta-Actin (Santa Cruz Biotechnology, sc-47778, 1:100), mouse anti-BACH1 (Abcam, ab128486, 1:2000), rabbit anti-PKNOX1 (ThermoFisher, PA5-30244, 1:750), mouse anti-GABPA (ThermoFisher, MA5-15419, 1:1000), goat anti-Rabbit-HRP (ThermoFisher, 31460, 1:5000) and goat anti-Mouse-HRP (ThermoFisher, 31430, 1:5000).

### snRNA-seq and analysis of graft tissue

Frozen grafts were processed and sequenced as described above for fetal tissue (10X-Multiome technology), or they were processed by Singleron and sequenced using the CeleScope scopeV3.0.1 (kit V2) technology (see Supplementary Table 6). Count matrices were generated as described for fetal tissue (cellranger-arc; v2.0.2; 10X Genomics), or using the CeleScope snRNA-seq mapping software version 2.0.7.

Quality control was performed as for fetal tissue with following modifications: As ATAC data was not analyzed here, only RNA-based filtering was performed to remove low quality cells. Cells with UMI counts < 500 or > 30,000, or with mitochondrial gene content > 2%, were excluded from the analysis. Datasets with <500 cells were removed, and large datasets were randomly subsampled to 2*median number of cells of the remaining samples, to reduce bias towards overrepresented samples. RNA data was then normalized by SCTransform and mapped onto the complete fetal dataset as reference (from Fig. 1; excluding non-cortical samples) using FindTransferAnchors(reference = seur_ref, query = seur, dims = 1:30, reference.reduction = “pca”) and MapQuery(anchorset = anchors, reference = seur_ref, query = seur, refdata = list(cluster_name = “cluster_name”, …), reference.reduction = “pca”, reduction.model = “umap”), to transfer metadata related to UMAP coordinates, cluster names, cell types, and developmental stage to the graft dataset. Cells mapping to the fetal excitatory lineage were re-mapped as above to the fetal excitatory lineage clusters (from Fig. 2a).

Differential cell abundance and gene expression analyses were performed with sccomp and DESeq2 using the transferred cluster labels, as described for fetal tissue. To correct for the use of a different sequencing technology (Singleron CeleScope) for some of the graft samples, DESeq2 analysis was performed with a log-likelihood-ratio test (LRT) including the sequencing technology as covariate, separately for each cluster, using the commands DESeqDataSetFromMatrix(counts_cluster, colData = meta_cluster, design = ∼seq_tech + group) and DESeq(test = “LRT”, reduced = ∼seq_tech), with a cutoff of padj <0.1.

For visualizing expression Z-scores including fetal and *in vitro* data, the combined bulk/pseudobulk count matrices were normalized using vst(), followed by batch correction with removeBatchEffect(batch = meta[[“seq_tech”]], group = meta$group) from the limma package^79^ (v3.58.1).

### Statistics and reproducibility

Data shown for representative experiments were repeated, with similar results, in at least two independent biological replicates and at least three technical replicates, unless otherwise noted.

## Acknowledgements

We thank the MRC-Wellcome Trust Human Developmental Biology Resource (HDBR), Dr Steven Lisgo and Dr Nita Solanski, the donors, and their families for the support, without which this study could not have been possible. Dr Aoife Murray and Dr Elizabeth Brockman for help with cell culture. Dr Phil Jun Kang and members of Prof Su Chun Zhang laboratory for help setting up the human iPSC-derived neuron culture protocols. Schung Hong Li and Dr Mynn Varela for help with the mouse colony maintenance. Samuela Pasculli for sharing analysis protocols for the qPCR experiments. Ms Amberlyn Tan for help with the RNA extraction and qPCR/WB. Dr Xavier Roca and Dr Huang Hua for assistance with the design and interpretation of the ASO experiments. Dr Joshua J. Gooley and Dr Xavier Roca, for comments on the manuscript.

## Author contributions

M.L. conceptualized and performed experiments with fetal tissue (stainings, sequencing), conceptualized and performed all sequencing data analyses, interpreted results, and wrote the manuscript. K.R. performed library preparations and sequencing, N.M. contributed to sequencing experiment design, Z.K. sourced and processed fetal tissue, performed stainings and imaging. M.S., S.S., W.L.T. and S.K.S. generated the neurons for the *in vitro* and *in vivo* experiments. K.H.U. and W.L.T. performed the qPCR, WB and ASO experiments and V.D.P., J.T., A.L. and B.A.V. the transplantation and graft extraction experiments. I.A. and D.N. provided the human iPSC models and derived neural progenitors used for transplantation experiments. B.P.L. contributed to study conceptualization and results interpretation. V.D.P. conceived and directed the study, acquired funding, interpreted results and wrote the manuscript.

## Data availability

Raw and processed sequencing data generated for this study will be made available under GEO accessions GSE305153. The fetal tissue dataset will also be available as interactive online resource on the CELLxGENE platform (https://cellxgene.cziscience.com/collections/0e9fd1d3-ef4c-47c6-a2e4-ef4bfadf7c79). Other data will be provided by the authors upon reasonable request.

## Code availability

The pipelines used for the analyses in this manuscript will be made available on Github (https://github.com/lattkem1/Down_Syndrome_Multiome).

